# Thanatotranscriptome: genes actively expressed after organismal death

**DOI:** 10.1101/058305

**Authors:** Alex E. Pozhitkov, Rafik Neme, Tomislav Domazet-Lošo, Brian G. Leroux, Shivani Soni, Diethard Tautz, Peter A. Noble

**Affiliations:** Department of Oral Health Sciences, University of Washington,Box 357444, Seattle, WA USA, 98195.; Max-Planck-Institute for Evolutionary Biology, August-Thienemann-Strasse 2, 24306 Ploen Germany.; Laboratory of Evolutionary Genetics, Division of Molecular Biology, Ruder Boškovic Institute, 10002 Zagreb, Croatia.; Catholic University of Croatia, Ilica 242, Zagreb, Croatia.; Department of Periodontics, School of Dentistry, University of Washington, Box 15 357444, Seattle, WA USA 98195.; Department of Biological Sciences, Alabama State University, Montgomery, AL, USA 36101-0271.; PhD Program in Microbiology, Alabama State University, Montgomery, AL, USA 36101-0271.

**Keywords:** Postmortem transcriptome, postmortem gene expression, Gene meters, calibrated DNA microarrays, thanatotranscriptome, cancer, signaling, development, immunity, apoptosis, transport, inflammation, epigenetic regulation, thermodynamic sink, transplantology, forensic science

## Abstract

A continuing enigma in the study of biological systems is what happens to highly ordered structures, far from equilibrium, when their regulatory systems suddenly become disabled. In life, genetic and epigenetic networks precisely coordinate the expression of genes -- but in death, it is not known if gene expression diminishes gradually or abruptly stops or if specific genes are involved. We investigated the unwinding of the clock by identifying upregulated genes, assessing their functions, and comparing their transcriptional profiles through postmortem time in two species, mouse and zebrafish. We found transcriptional abundance profiles of 1,063 genes were significantly changed after death of healthy adult animals in a time series spanning from life to 48 or 96 h postmortem. Ordination plots revealed non-random patterns in profiles by time. While most thanatotranscriptome (thanatos-, Greek *defn*. death) transcript levels increased within 0.5 h postmortem, some increased only at 24 and 48 h. Functional characterization of the most abundant transcripts revealed the following categories: stress, immunity, inflammation, apoptosis, transport, development, epigenetic regulation, and cancer. The increase of transcript abundance was presumably due to thermodynamic and kinetic controls encountered such as the activation of epigenetic modification genes responsible for unraveling the nucleosomes, which enabled transcription of previously silenced genes (e.g., development genes). The fact that new molecules were synthesized at 48 to 96 h postmortem suggests sufficient energy and resources to maintain self-organizing processes. A step-wise shutdown occurs in organismal death that is manifested by the apparent upregulation of genes with various abundance maxima and durations. The results are of significance to transplantology and molecular biology.

## INTRODUCTION

A healthy adult vertebrate is a complex biological system capable of highly elaborate functions such as the ability to move, communicate, and sense the environment -- all at the same time. These functions are tightly regulated by genetic and epigenetic networks through multiple feedback loops that precisely coordinate the expression of thousands of genes at the right time, in the right place, and in the right level [1]. Together, these networks maintain homeostasis and thus sustain ‘life’ of a biological system.

While much is known about gene expression circuits in life, there is a paucity of information about what happens to these circuits after organismal death. For example, it is not well known whether gene expression diminishes gradually or abruptly stops in death -- nor whether specific genes are newly expressed or upregulated. In organismal ‘death’, defined here as the cessation of the highly elaborate system functions in vertebrates, we conjecture that there is a gradual disengagement and loss of global regulatory networks, but this could result in a regulatory response of genes involved in survival and stress compensation. To test this, we examined postmortem gene expression in two model organisms: the zebrafish, *Danio rerio*, and the house mouse, *Mus musculus*. The purpose of the research was to investigate the “unwinding of the clock” by identifying genes whose expression increases (i.e., nominally upregulated) and assessing their functions based on the primary literature. The biological systems investigated in this study are different from those examined in other studies, such as individual dead and/or injured cells in live organisms, i.e., apoptosis and necrosis (reviewed in refs. [2–5]). In contrast to previous studies, gene expression from the entire *D. rerio* body, and the brains and livers of *M. musculus* were assessed through postmortem time. Gene expression was measured using the ‘Gene Meter’ approach that precisely reports gene transcript abundances based on a calibration curve for each microarray probe [6].

## MATERIALS AND METHODS

**Induced death and postmortem incubation.** <u>Zebrafish</u> Forty-four female *Danio rerio* were transferred from several flow-through aquaria kept at 28°C to a glass beaker containing 1 L of aquarium water. Four individuals were immediately taken out, snap frozen in liquid nitrogen, and stored in Falcon tubes at −80°C (two zebrafish per tube). These samples were designated as the first set of live controls. A second set of live controls was immersed in an open cylinder (described below). Two sets of live controls were used to determine if putting the zebrafish back into their native environment had any effects on gene expression (we later discovered no significant effects).

The rest of the zebrafish were subjected to sudden death by immersion in a “kill” chamber. The chamber consisted of an 8 L styrofoam container filled with chilled ice water. To synchronize the death of the rest of the zebrafish, they were transferred to an open cylinder with a mesh-covered bottom and the cylinder was immersed into the kill chamber. After 20 to 30 s of immersion, four zebrafish were retrieved from the chamber, snap frozen in liquid nitrogen, and stored at −80°C (two zebrafish per Falcon tube). These samples were designated as the second set of live controls. The remaining zebrafish were kept in the kill chamber for 5 min and then the cylinder was transferred to a flow-through aquarium kept at 28°C so that they were returned to their native environment.

Postmortem sampling of the zebrafish occurred at: time 0, 15 min, 30 min, 1 h, 4 h, 8 h, 12 h, 24 h, 48 h, and 96 h. For each sampling time, four expired zebrafish were retrieved from the cylinder, snap frozen in liquid nitrogen, and stored at -80°C in Falcon tubes (two zebrafish to a tube). One zebrafish sample was lost, but extraction volumes were adjusted to one organism.

<u>Mouse</u> The mouse strain C57BL/6JRj (Janvier SAS, France) was used for our experiments. The mice were 20-week old males of approximately the same weight. The mice were highly inbred and were expected to have a homogenous genetic background. Prior to euthanasia, the mice were kept at room temperature and were given *ad libitum* access to food and water. Each mouse was euthanized by cervical dislocation and placed in an individual plastic bag with holes to allow air / gas exchange. The bagged carcasses were kept at room temperature in a large, open polystyrene container. Sampling of the deceased mice began at 0 h (postmortem time zero) and continued at 30 min, 1 h, 6 h, 12 h, 24 h and 48 h postmortem. At each sample time, 3 mice were sampled (except for 48h where 2 mice were sampled) and the entire brain (plus stem) and two portions of the liver were extracted from each mouse. For liver samples, clippings were taken from the foremost and rightmost lobes of the liver. The brain and liver samples were snap frozen in liquid nitrogen and stored individually in Falcon tubes at −80°C.

The euthanasia methods outlined above are approved by the American Veterinary Medical Association (AVMA) Guidelines for Euthanasia (www.avma.org) and carried out by personnel of the Max-Planck-Institute for Evolutionary Biology (Ploen, Germany). All animal work: followed the legal requirements, was registered under number V312- 72241.123-34 (97-8/07) and approved by the ethics commission of the Ministerium fur Landwirtschaft, Umwelt und landliche Raume, Kiel (Germany) on 27. 12. 2007.

**RNA extraction, labeling, hybridization and DNA microarrays.** The number of biologically distinct organisms was 43 for zebrafish and 20 for mice. Samples from two fish were pooled for analysis, resulting in two replicate measurements at each time point. The number of replicated measurements for mice was three at each of the first six time points and two at 48h. Thus, the total number of samples analyzed was 22 for zebrafish and 20 for mice. For the zebrafish, samples were mixed with 20 ml of Trizol and homogenized using a TissueLyzer (Qiagen). For the mice, 100 mg of brain or liver samples were mixed with 1 ml of Trizol and homogenized. One ml of the emulsion from each sample was put into a fresh 1.5 ml centrifuge tube for RNA extraction and the rest was frozen at −80°C.

RNA was extracted by adding 200 μl of chloroform, vortexing the sample, and incubating it at 25°C for 3 min. After centrifugation (15 min at 12000 × g at 4°C), the supernatant (approx. 350 μl) was transferred to a fresh 1.5 ml tube containing an equal volume of 70% ethanol. The tube was vortexed, centrifuged and purified following the procedures outlined in the PureLink RNA Mini Kit (Life Technologies, USA).

The isolated RNA, 400 ng per sample, was labeled, purified and hybridized according to the One-Color Microarray-based Gene Expression Analysis (Quick Amp Labeling) with Tecan HS Pro Hybridization kit (Agilent Technologies). For the zebrafish, the labeled RNA was hybridized to the Zebrafish (v2) Gene Expression Microarray (Design ID 019161). For the mouse, the labeled RNA was hybridized to the SurePrint G3 Mouse GE 8x60K Microarray Design ID 028005 (Agilent Technologies). The microarrays were loaded with 1.65 μg of labeled cRNA for each postmortem sample.

**Microarray calibration.** Oligonucleotide (60 nt) probes on the zebrafish and mouse microarrays were calibrated using pooled labeled cRNA of all zebrafish and all mouse postmortem samples, respectively. The dilution series for the Zebrafish array was created using the following concentrations of labeled cRNA: 0.41, 0.83, 1.66, 1.66, 1.66, 3.29, 6.60, and 8.26 μg. The dilution series for the Mouse arrays was created using the following concentrations of labeled cRNA: 0.17, 0.33, 0.66, 1.32, 2.64, 5.28, 7.92, and 10.40 μg. Calibration involved plotting the signal intensities of the probes against a dilution factor and determining the isotherm model (e.g., Freundlich and/or Langmuir) that best fit the relationship between signal intensities and gene abundances.

Consider zebrafish gene transcripts targeted by A_15_P110618 (which happens to be one of the transcriptional profiles of gene *Hsp70.3* shown in Fig 2A). External file FishProbesParameters.txt shows that a Freundlich model best fit the dilution curve with R^2^=0.99. The equation for this probe is the following: 
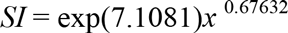
where *SI* is the observed average signal intensity for dilution *x*. The gene abundance *G* was calculated by inverting this equation. For this probe, signal intensity at each postmortem time, *SI_t_*, is determined by the equation: *G*= (*SI_t_*/ exp(7.1081))^(1/0.67632). Specifically, consider two biological replicates of 15 min postmortem zebrafish, the signal intensities of the probe A_15_P110618 are 770.5 and 576.0, which translates into the abundances 0.50 and 0.33 arbitrary units (a.u.) respectively. The target abundances were further converted to log10 and are shown in external file Fish_log10_AllProfiles.txt.

Details of the calibration protocols to calculate gene expression values, i.e., mRNA relative abundances, are provided in our recent paper where we describe the “Gene Meter” [6].

**Statistical analysis.** Abundance levels were log-transformed for analysis to stabilize the variance. A one-sided Dunnett’s T-statistic was applied to test for increase at one or more postmortem times compared to live control (fish) or time 0 (mouse). A bootstrap procedure with 10^9^ simulations was used to determine the critical value for the Dunnett statistics in order to accommodate departures from parametric assumptions and to account for multiplicity of testing. The profiles for each gene were centered by subtracting the mean values at each postmortem time point to create “null” profiles. Bootstrap samples of the null profiles were generated to determine the 95th percentile of the maximum (over all genes) of the Dunnett statistics. Significant postmortem upregulated genes were selected as those having Dunnett T values larger than the 95th percentile. Only significantly upregulated genes were retained for further analyses.

Orthogonal transformation of the abundances to their principal components (PC) was conducted and the results were graphed on a 2 dimensional ordination plot. The *m* × *n* matrix of abundances (sampling times by number of gene transcripts), which is 10 × 548 for zebrafish and 7 × 515 for mouse, was used to produce an *m* × *m* matrix D of Euclidean distances between all pairs of sampling times. Principal component analysis (PCA) was performed on the matrix of distances, D. To investigate and visualize differences between the sampling times, a scatterplot of the first two principal components (PC1 and PC2) was created. To establish relative contributions of the gene transcripts, the projection of each sampling time onto the (PC1, PC2) plane was calculated and those genes with high correlations (>=0.70) between abundances and either component (PC1 or PC2) were displayed as a biplot.

**Gene annotation and functional categorization.** Microarray probe sequences were individually annotated by performing a BLASTN search of the zebrafish and mouse NCBI databases (February, 2015). The gene annotations were retained if the bit score was greater than or equal to 100 and the annotations were in the correct 5’ to 3’ orientation. Transcription factors, transcriptional regulators, and cell signaling components (e.g., receptors, enzymes, and messengers) were identified as global regulatory genes. The rest were considered response genes.

Functional categorizations were performed by querying the annotated gene transcripts in the primary literature and using UniProt (www.uniprot.org). Genes not functionally categorized to their native organism (zebrafish or mouse) were categorized to genes of phylogenetically related organisms (e.g., human). Cancer-related genes were identified using a previously constructed database (see Additional File 1: Table S1 in [7]).

## RESULTS

Similar quantities of total mRNA were extracted from zebrafish samples for the first 12 h postmortem (avg. 1551 ng per μl tissue extract) then the quantities abruptly decreased with postmortem time (Table S1). The quantities of total mRNA extracted from the mouse liver samples were about the same for the first 12 h postmortem (avg. of 553 ng per μl tissue extract) then they increased with time (Table S2). The quantities of total mRNA extracted from the mouse brain samples were similar (avg. of 287 ng per μl tissue extract) for all postmortem times (Table S2). Hence, the amount of total mRNA extracted depended of the organism/organ/tissue and time.

Calibration of the microarray probes and determination of the transcript abundances at each postmortem sampling time produced a fine-grain series of data for the zebrafish and the mouse. Approximately 84.3% (36,811 of 43,663) zebrafish probes and 67.1% (37,368 of 55,681) mouse probes were found to provide a suitable dose-response curve for calibration (data available upon request).

Figure 1 shows the sum of all gene abundances calculated from the calibrated probes with postmortem time. In general, the sum of all abundances decreased with time, which means that less targets hybridized to the microarray probes. In the zebrafish, mRNA decreased abruptly at 12 h postmortem (Fig 1A), while for the mouse brain Fig 1B), total mRNA increased in the first hour and then gradually decreased. For the mouse liver, mRNA gradually decreased with postmortem time. The fact that total mRNA shown in Figs 1A and 1B mirrors the electrophoresis banding patterns shown in (Fig S1 and S2 (ignoring the 28S and 18S rRNA bands) indicates a general agreement of the Gene Meter approach to another molecular approach (i.e., Agilent Bioanalyzer). Hence, total mRNA abundances depended on the organism (zebrafish, mouse), organ (brain, liver), and postmortem time, which are aligned with previous studies [8,9,10,11,12,13].

**Fig 1.**
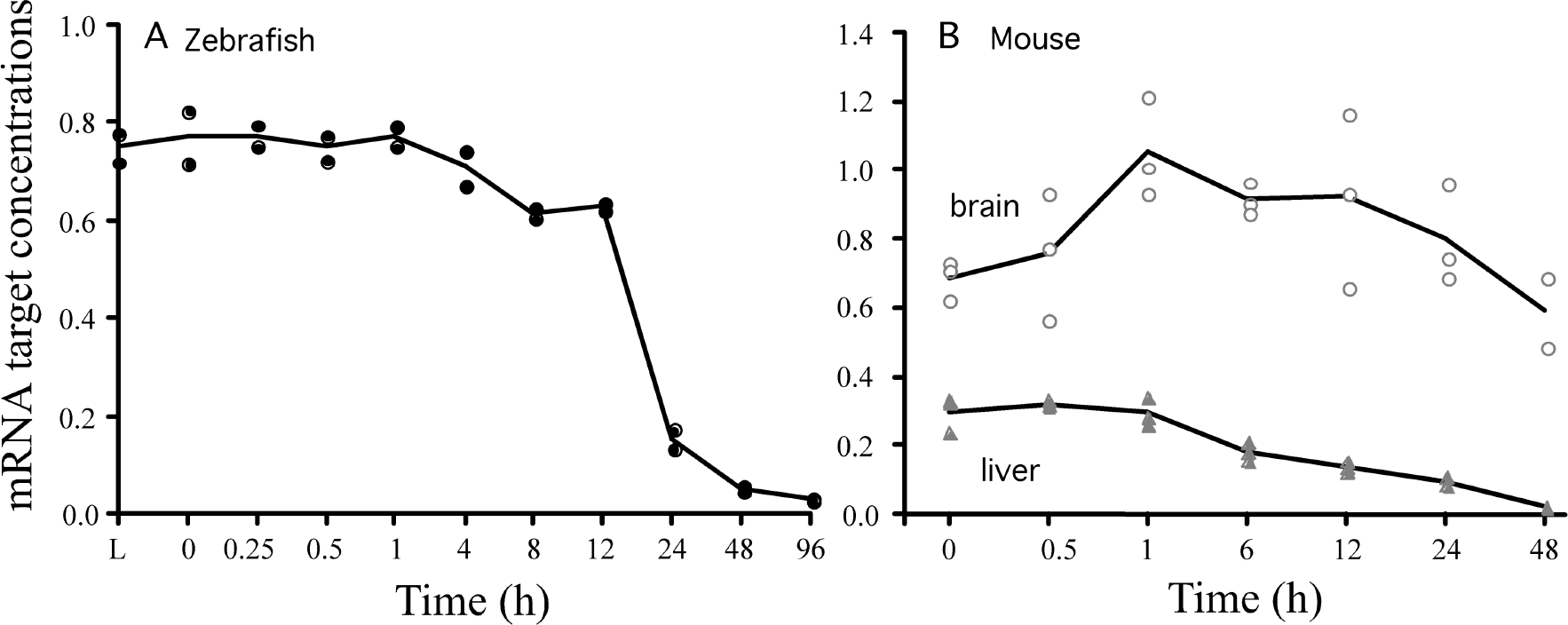
**Total mRNA abundance (arbitrary units, a.u.) by postmortem time determined using all calibrated microarray probes. A, extracted from whole zebrafish; B, extracted from brain and liver tissues of whole mice. Each datum point represents the mRNA from two organisms in the zebrafish and a single organism in the mouse**.

The abundance of a gene transcript is determined by its rate of synthesis and its rate of degradation [14]. We focused on genes that show a significant increase in RNA abundance -- relative to live controls -- because these genes are likely to be actively transcribed in organismal death despite an overall decrease in total mRNA with time. An upregulated transcription profile was defined as one having at least one time point where the abundance was statistically higher than that of the control (Fig 2 A to 2C). It is important to understand that the entire profiles, i.e., 22 data points for the zebrafish and 20 points for the mouse, were subjected to a statistical test to determine significance (see Materials and Methods). We found 548 zebrafish transcriptional profiles and 515 mouse profiles were significantly upregulated. The fact that there are upregulated genes is consistent with the notion that there is still sufficient energy and functional cellular machinery for transcription to occur -- long after organismal death.

**Fig 2.**
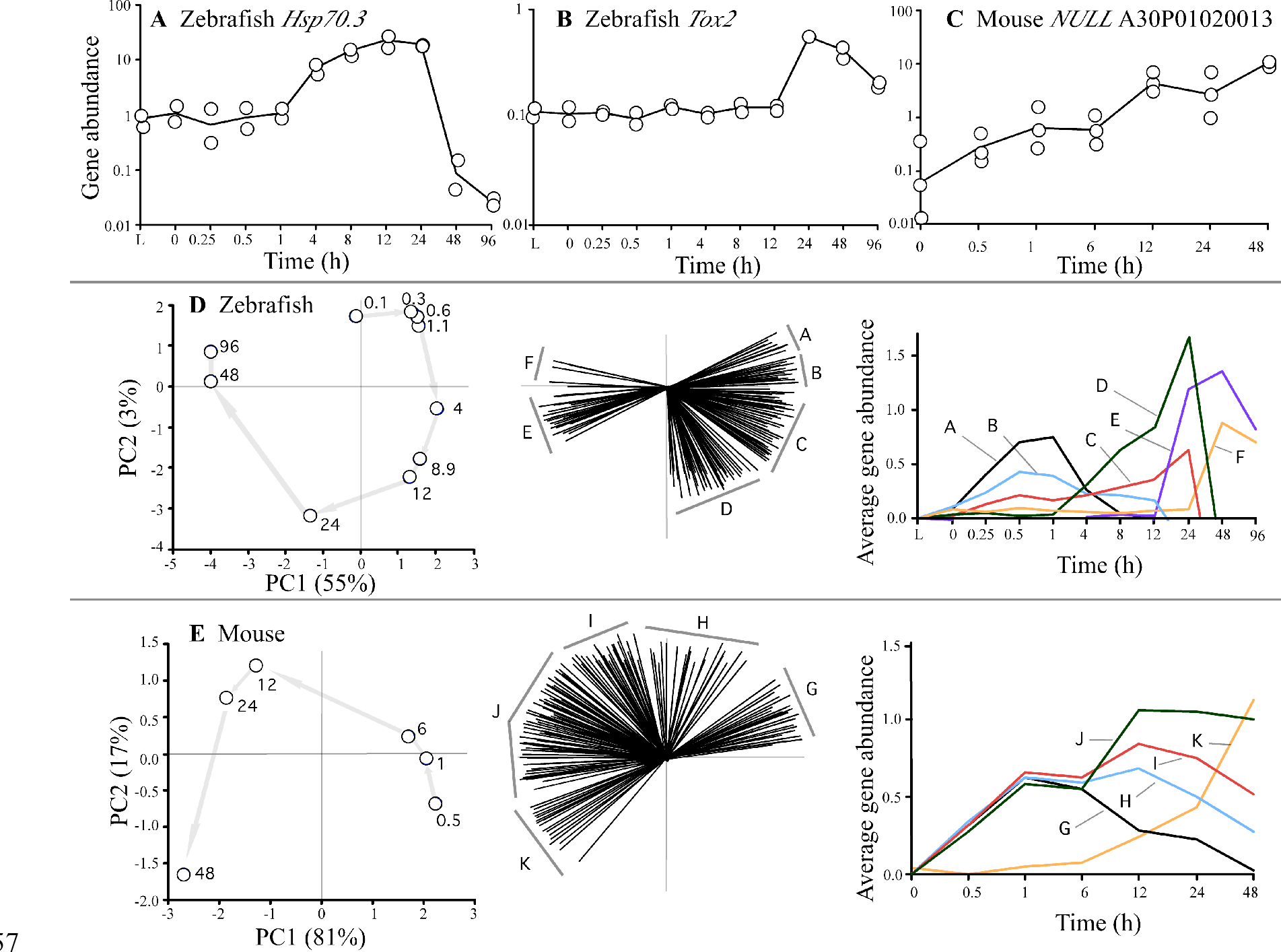
**Transcriptional profiles of representative genes (a.u.), ordination plots based on transcript abundances by postmortem time (h) with corresponding transcript contributions (biplots), and averaged transcript abundances by group. (A) Transcriptional profile of the *Hsp70.3* gene, (B) the *Tox2* gene, and (C) the NULL (i.e., no annotation, probe number shown) gene as a function of postmortem time. Each datum point was derived from the mRNA of two zebrafish or one mouse. (D) Ordination plots of the zebrafish and (E) mouse were based on all upregulated gene profiles by postmortem time (h). Gene transcripts in the biplots were arbitrarily assigned alphabetical groups based on their positions in the ordination. The average transcript abundances for each group are shown**.

Based on GenBank gene annotations, we found that among the upregulated genes for the zebrafish, 291 were protein-coding genes (53%) and 257 non-coding mRNA (47%) and, for the mouse, 324 known protein-coding genes (63%), 190 non-coding mRNA (37%), and one control sequence of unknown composition. Hence, about 58% of the total upregulated genes in the zebrafish and mouse are known and the rest (42%) are noncoding RNA.

Examples of genes yielding transcripts that significantly increased in abundance with postmortem time are: the ‘Heat shock protein’ (*Hsp70.3*) gene, the ‘Thymocyte selection-associated high mobility group box 2’ (*Tox2*) gene, and an unknown (*NULL*) gene (Fig 2A to 2C). While the *Hsp70.3* transcript increased after 1 h postmortem to reach a maximum at 12 h, the *Tox2* transcript increased after 12 h postmortem to reach a maximum at 24 h, and the *NULL* transcript consistently increased with postmortem time. These figures provide typical examples of transcript profiles and depict the high reproducibility of the sample replicates as well as the quality of output obtained using the Gene Meter approach.

### Non-random patterns in transcript profiles

Ordination plots of the significantly upregulated transcript profiles revealed prominent differences with postmortem time (Fig 2D and 2E), suggesting the expression of genes followed a discernible (non-random) pattern in both organisms. The biplots showed that 203 zebrafish transcript profiles and 226 mouse profiles significantly contributed to the ordinations. To identify patterns in the transcript profiles, we assigned them to groups based on their position in the biplots. Six profile groups were assigned for the zebrafish (A to F) and five groups (G to K) were assigned for the mouse. Determination of the average gene transcript abundances by group revealed differences in the shapes of the averaged profiles, particularly the timing and magnitude of peak transcript abundances, which accounted for the positioning of data points in the ordinations.

Genes coding for global regulatory functions were examined separately from others (i.e., response genes). Combined results show that about 33% of the upregulated genes in the ordination plots were involved in global regulation with 14% of these encoding transcription factors/transcriptional regulators and 19% encoding cell signaling proteins such as enzymes, messengers, and receptors (Table S3). The response genes accounted for 67% of the upregulated transcripts.

The genes were assigned to 22 categories (File S8) with some genes having multiple categorizations. For example, the Eukaryotic Translation Initiation Factor 3 Subunit J-B (*Eif3j2*) gene was assigned to protein synthesis and cancer categories [15].

Genes in the following functional categories were investigated: stress, immunity, inflammation, apoptosis, solute/ion/protein transport, embryonic development, epigenetic regulation and cancer. We focused on these categories because they were common to both organisms and contained multiple genes, and they might provide explanations for postmortem upregulation of genes (e.g., epigenetic gene regulation, embryonic development, cancer). The transcriptional profiles of the genes were plotted by category and each profile was ordered by the timing of the upregulation and peak transcript abundance. This allowed comparisons of gene expression dynamics as a function of postmortem time for both organisms. For each category, we provided the gene name and function and compared expression dynamics within and between the organisms.

### Stress response

In organismal death, we anticipated the upregulation of stress response genes because these genes are activated in life to cope with perturbations and to recover homeostasis [16]. The stress response genes were assigned to three groups: heat shock protein (*Hsp*), hypoxia-related, and ‘other’ responses such as oxidative stress.

***Hsp*** In the zebrafish, upregulated *Hsp* genes included: ‘Translocated promoter region’ (*Tpr*), *Hsp70.3*, and *Hsp90* (Fig 3). The *Tpr* gene encodes a protein that facilitates the export of the *Hsp* mRNA across the nuclear membrane [17] and has been implicated in chromatin organization, regulation of transcription, mitosis [18] and controlling cellular senescence [19]. The *Hsp70.3* and *Hsp90* genes encode proteins that control the level of intracellular calcium [20], assist with protein folding, and aid in protein degradation [21].

In the mouse, upregulated *Hsp* genes included: *Tpr*, Hsp-associated methyltransferase (*Mettl21*) and Heat Shock Protein 1 (*Hspel*) (Fig 3). The *Mettl21* gene encodes a protein modulating *Hsp* functions [22]. The *Hspel* gene encodes a chaperonin protein that assists with protein folding in the mitochondria [23].

The timing and duration of *Hsp* upregulation varied by organism. In general, activation of *Hsp* genes occurred much later in the zebrafish than the mouse (4 h vs. 0.5 h postmortem, respectively). There were also differences in transcript abundance maxima since, in the zebrafish, maxima were reached at 9 to 24 h, while in the mouse maxima were reached at 12 to 24 h. Previous studies have examined the upregulation of *Hsp70.3* with time in live serum-stimulated human cell lines [24]. In both the zebrafish and human cell lines the *Hsp70.3* gene transcript reached maximum abundance at about 12 h (Fig 2A), indicating the same reactions are occurring in life and death.

**Fig 3.**
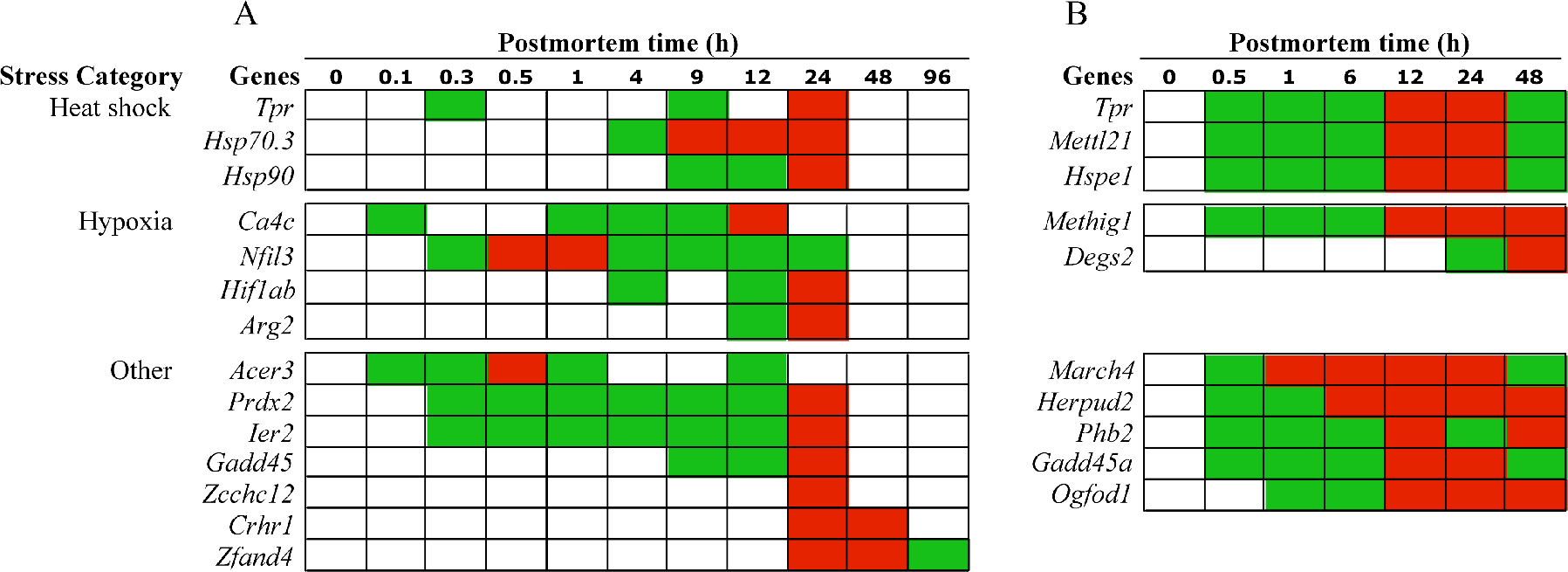
**Upregulated stress response genes by postmortem time (h) and stress category. A. zebrafish; B, mouse. Green, intermediate value; Red, maximum value**.

#### Hypoxia

In the zebrafish, upregulated hypoxia-related genes included: Carbonic anhydrase 4 (*Ca4c*), Nuclear factor interleukin-3 (*Nfil3*), Hypoxia-inducible factor 1-alpha (*Hiflab*) and Arginase-2 (*Arg2*) (Fig 3). The Carbonic anhydrase 4 (*Ca4c*) gene encodes an enzyme that converts carbon dioxide into bicarbonate in response to anoxic conditions [25]. The *Nfil3* gene encodes a protein that suppresses hypoxia-induced apoptosis [26] and activates immune responses [27]. The *Hiflab* gene encodes a transcription factor that prepares cells for decreased oxygen [28]. The *Arg2* gene encodes an enzyme that catalyzes the conversion of arginine to urea under hypoxic conditions [29]. Of note, the accumulation of urea presumably triggered the upregulation of the *Slc14a2* gene at 24 h, reported in the Transport Section (below).

In the mouse, upregulated hypoxia genes included: Methyltransferase hypoxia inducible domain (*Methigl*) and Sphingolipid delta-desaturase (*Degs2*) (Fig 3). The *Methigl* gene encodes methyltransferase that presumably is involved in gene regulation [30]. The *Degs2* gene encodes a protein that acts as an oxygen sensor and regulates ceramide metabolism [31]. Ceramides are waxy lipid molecules in cellular membranes that regulate cell growth, death, senescence, adhesion, migration, inflammation, angiogenesis and intracellular trafficking [32].

The activation of the *Ca4c* gene in the zebrafish indicates a build up of carbon dioxide at 0. 1 to 1 h postmortem in the zebrafish presumably due lack of blood circulation. The upregulation of the *Nfil3* gene in the zebrafish and *Methig1* gene in the mouse suggests hypoxic conditions existed within 0.5 h postmortem in both organisms. The upregulation of other hypoxia genes varied with postmortem time, with the upregulation of *Hiflab, Arg2*, and *Degs2* genes occurring at 4 h, 12 h and 24, respectively.

#### Other stress responses

In the zebrafish, upregulated response genes included: Alkaline ceramidase 3 (*Acer3*), Peroxirodoxin 2 (*Prdx2*), Immediate early (*Ier2*), Growth arrest and DNA-damage-inducible protein (*Gadd45a*), Zinc finger CCH domain containing 12 (Zcchc12), Corticotropin releasing hormone receptor 1 (*Crhr1*), and Zinc finger AN1-type domain 4 (*Zfand4*) (Fig 3). The *Acer3* gene encodes a stress sensor protein that mediates cell-growth arrest and apoptosis [33]. The *Prdx2* gene encodes an antioxidant enzyme that controls peroxide levels in cells [34] and triggers production of *Tnfa* proteins that induce inflammation [35]. The *Ier2* gene encodes a transcription factor involved in stress response [36]. The *Gadd45a* gene encodes a stress protein sensor that stops the cell cycle [37], modulates cell death and survival, and is part of the signaling networks in immune cells [38]. The *Zcchc12* gene encodes a protein involved in stress response in the brain [39]. The *Crhr1* and *Zfand4* genes encode stress proteins [40,41].

While the *Acer3, Prdx2*, and *Ier2* genes were activated within 0.3 h postmortem, indicating a changed physiological state; the *Gadd45a* gene was activated at 9 h and the other genes (*Zcchc12, Crhr1, Zfand4*) were activated at 24 h postmortem.

In the mouse, upregulated stress response genes included: Membrane-associated RING-CH 4 (*March4*), Homocysteine-responsive endoplasmic reticulum-resident ubiquitin-like domain member 2 (*Herpud2*), Prohibitin-2 (*Phb2*), *Gadd45a*, and Two-oxoglutarate and iron-dependent oxygenase domain-containing 1 (*Ogfod1*) (Fig 3). The *March4* gene encodes an immunologically-active stress response protein [42]. The *Herpud2* gene encodes a protein that senses the accumulation of unfolded proteins in the endoplasmic reticulum [43]. The *Phb2* gene encodes a cell surface receptor that responds to mitochondrial stress [44]. The *Ogfod1* gene encodes a stress-sensing protein [45].

Note that the stress genes in the mouse were all activated within 1 h postmortem and remained upregulated for 48 h.

#### Summary of stress response

In both organisms, organismal death activated heat shock, hypoxia, and other stress genes, which varied in the timing and duration of upregulation within and between organisms. Consider, for example, the *Tpr* and *Gadd45a* genes, which were common to both organisms. While the *Tpr* genes were upregulated within 0.5 h postmortem in both organisms, the *Gadd45a* gene was upregulated at 9 h in the zebrafish and 0.5 h in the mouse. In addition, the transcription profile of the *Tpr* gene was more variable in the zebrafish than the mouse since it was activated at 0.3 h, 9 h and 24 h postmortem, which suggest the gene might be regulated through a feedback loop. In contrast, in the mouse, the *Tpr* gene was upregulated at 0.5 h and the transcripts reached peak abundance at 12 and 24 h postmortem.

Taken together, the stress genes were activated in both organisms to compensate for a loss in homeostasis.

### Innate and adaptive immune responses

In organismal death, we anticipated the upregulation of immune response genes because vertebrates have evolved ways to protect the host against infection in life, even under absolutely sterile conditions [46]. Inflammation genes were excluded from this section (even though they are innate immune genes) because we examined them in a separate section (below).

In the zebrafish, upregulated immunity genes included: Early growth response-1 and -2 (*Egr1, Egr2*), Interleukin-1b (*Il1b*), L-amino acid oxidase (*Laao*), Interleukin-17c (*Il17c*), Membrane-spanning 4-domains subfamily A member 17A.1 (*Ms4a17.a1*), Mucin-2 (*Muc2*), Immunoresponsive gene 1 (*Irg1*), Interleukin- 22 (Il22), Ubl carboxyl-terminal hydrolase 18 (*Usp18*), ATF-like 3 (*Batf3*), Cytochrome b-245 light chain (*Cyba*), and ‘Thymocyte selection-associated high mobility group” box protein family member 2 (*Tox2*) (Fig 4). The *Egr1* and *Egr2* genes encode proteins that regulate B and T cell functions in adaptive immunity [47,48]. The *Il1b* gene encodes an interleukin that kills bacterial cells through the recruitment of other antimicrobial molecules [49]. The *Laao* gene encodes an oxidase involved in innate immunity [50]. The *Il17c* and *Il22* genes encode interleukins that work synergically to produce antibacterial peptides [51]. The *Ms4a17.a1* gene encodes a protein involved in adaptive immunity [52]. The*Muc2* gene encodes a protein that protects the intestinal epithelium from pathogenic bacteria [53]. The *Irg1* gene encodes an enzyme that produces itaconic acid, which has antimicrobial properties [54]. The *Usp18* gene encodes a protease that plays a role in adaptive immunity [55]. The *Batf3* gene encodes a transcription factor that activates genes involved in adaptive immunity [56]. The *Cyba* gene encodes an oxidase that is used to kill microorganisms [57]. The *Tox2* gene encodes a transcription factor that regulates Natural Killer (NK) cells of the innate immune system [58].

**Fig 4.**
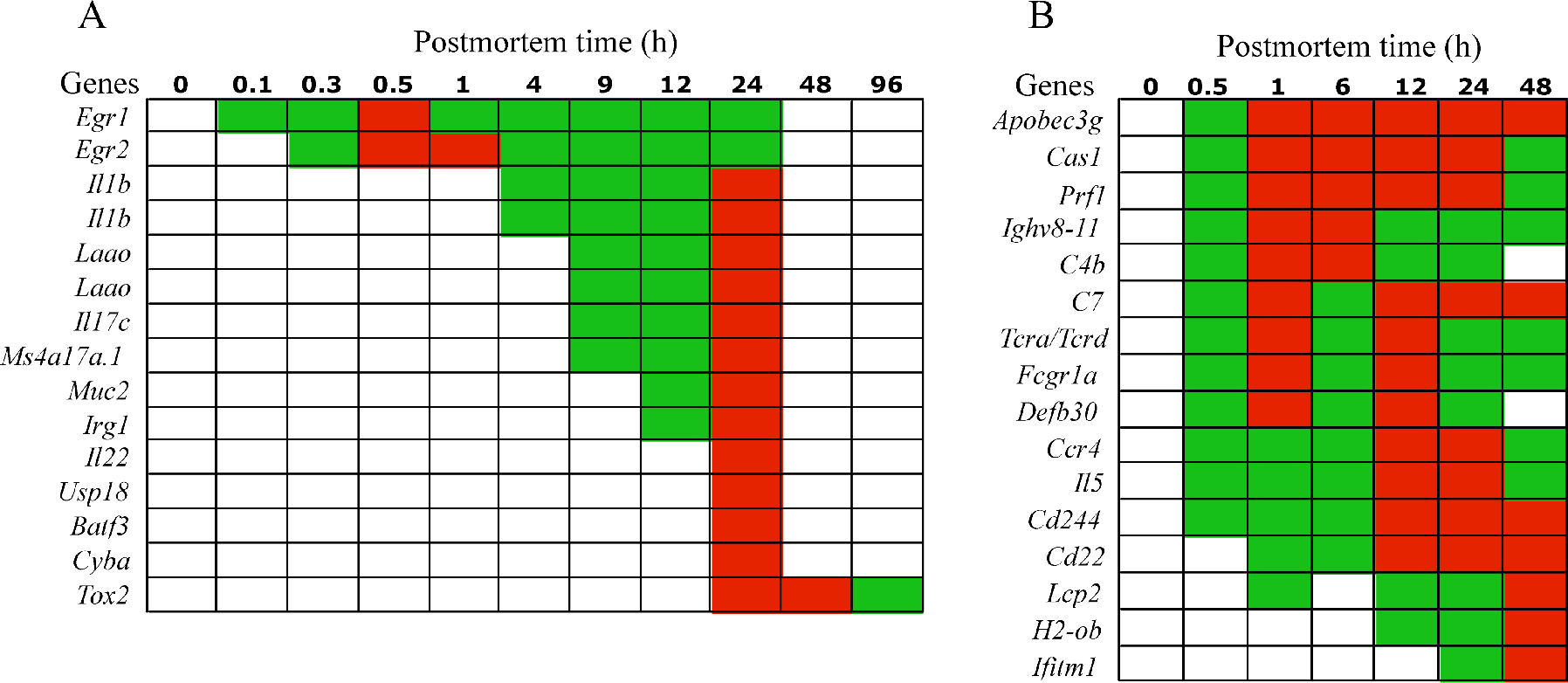
**Upregulated immunity genes by postmortem time (h). A. zebrafish; B, mouse. Green, intermediate value; Red, maximum value. Some transcripts were represented by two different probes (e.g. *Illb*, *Laao*)**.

Upregulation of immunity genes in the zebrafish occurred at different times with varying durations. While genes involved in adaptive immunity were upregulated at 0.1 to 0.3 h (*Egr*), 9 h (*Ms4a17.a1*) and 24 h (*Usp18, Batf3*) postmortem, genes involved in innate immunity were upregulated at 4 h (*Il1b*), 9 h (*Laao, Il17c*), 12 h (*Muc2, Irg1*) and 24 h (*Il22, Cyba, Tox2*) indicating a multi-pronged and progressive approach to deal with injury and the potential of microbial invasion.

In the mouse, upregulated antimicrobial genes included: Catalytic polypeptide-like 3G (*Apobec3g*), CRISPR-associated endonuclease (*Cas1*), Perforin-1 (*Prf1*), Immunoglobulin heavy variable 8-11 (*Ighv8-11*), C4b-binding protein (C4b), Complement component C7 (C7), T cell receptor alpha and delta chain (*Tcra/Tcrd*), High affinity immunoglobulin gamma Fc receptor I (*Fcgr1a*), Defensin (*Defb30*), Chemokine- 4 (Ccr4), Interleukin-5 (*Il5*), NK cell receptor 2B4 (*Cd244*), Cluster of differentiation-22 (Cd22), Lymphocyte cytosolic protein 2 (*Lcp2*), Histocompatibility 2 O region beta locus (*H2ob*) and Interferon-induced transmembrane protein 1 (*Iftm1*) (Fig 4). The *Apobec3g* gene encodes a protein that plays a role in innate anti-viral immunity [59]. The *Cas1* gene encodes a protein involved in regulating the activation of immune systems [60,61,62,63]. The *Prf1, C7*, and *Defb30* genes encode proteins that kill bacteria by forming pores in the plasma membrane of target cells [64,65,66]. The *Ighv8-11* gene encodes an immunoglobulin of uncertain function. The *C4b* gene encodes a protein involved in the complement system [67]. The *Tcra/Tcrd* genes encode proteins that play a role in the immune response [68]. The *Fcgr1a* gene encodes a protein involved in both innate and adaptive immune responses [69]. The *Ccr4* gene encodes a cytokine that attracts leukocytes to sites of infection [70]. The *Il5* gene encodes an interleukin involved in both innate and adaptive immunity [71,72]. The *Cd244* and *Cd22* genes encode proteins involved in innate immunity [73]. The *Lcp2* gene encodes a signal-transducing adaptor protein involved in T cell development and activation [74]. The *H2ob* gene encodes a protein involved in adaptive immunity. The *Ifitml* gene encodes a protein that has antiviral properties [75].

Most immune response genes were upregulated within 1 h postmortem in the mouse (*n*=14 out of 16 genes), indicating a more rapid response than that of the zebrafish.

#### Summary of immune response

The upregulated immune response genes in both organisms included innate and adaptive immunity components. An interesting phenomenon observed in the mouse (but not zebrafish) was that four genes (C7, *Tcra/Tcrd, Fcgrla*, and *Defb30*) reached maximum transcript abundance at two different postmortem times (i.e., 1 h and 12 h) while others reached only one maximum. The variability in the gene transcript profiles suggests their regulation is effected by feedback loops.

#### Inflammation response

We would anticipate the upregulation of inflammation genes in organismal death because inflammation is an innate immunity response to injury. In the zebrafish, upregulated inflammation genes included: *Egr1, Egr2, Il1b*, Tumor necrosis factor receptor (*Tnfrsf19*), Heme oxygenase 1 (*Hmox1*), Tumor necrosis factor (*Tnf*), G-protein receptor (*Gpr31*), Interleukin-8 (Il8), Tumor necrosis factor alpha (*Tnfa*), Nuclear factor (NF) kappa B (*Nfkbiaa*), MAP kinase-interacting serine/threonine kinase 2b (*Mknk2b*), and Corticotropin-releasing factor receptor 1 (*Crhr1*) (Fig 5). The *Egr1* and *Egr2* genes encode transcription factors that are pro- and anti- inflammatory, respectively [47,48,76]. The *Il1b* gene encodes a pro-inflammatory cytokine that plays a key role in sterile inflammation [77,78]. The *Tnfrsf19* gene encodes a receptor that has pro-inflammatory functions [79]. The *Hmox1* gene encodes an enzyme that has anti-inflammatory functions and is involved in Heme catabolism [80,81]. The *Tnf* and *Tnfa* genes encode pro-inflammatory proteins. The *Gpr31* gene encodes a pro-inflammatory protein that activates the NF-kB signaling pathway [82]. The *Il8* gene encodes a cytokine that has pro-inflammatory properties [83]. The *Nfkbiaa* gene encodes a protein that integrates multiple inflammatory signaling pathways including *Tnf* genes [84]. The *Mknk2b* gene encodes a protein kinase that directs cellular responses and is pro-inflammatory [85]. The *Crhr1* gene modulates anti-inflammatory responses [86].

**Fig 5.**
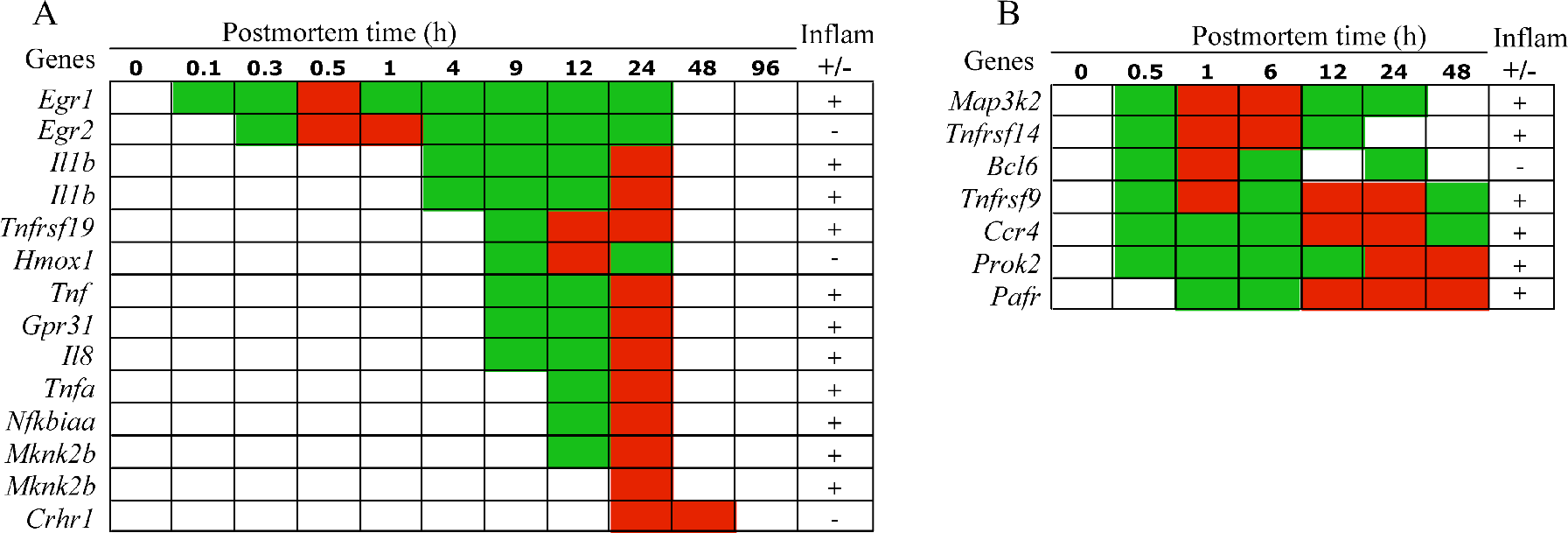
**Upregulated inflammation genes by postmortem time (h). A. zebrafish; B, mouse. Inflammation, Pro-, +; Anti, − . Green, intermediate value; Red, maximum value. The *Illb* and *Mknk2b* genes were represented by two different probes**.

The upregulation of pro-inflammatory *Egrl* gene at 0.1 h was followed by the upregulation of anti-inflammatory *Egr2* gene at 0.2 h, suggesting the transcription of one gene is affecting the regulation of another (Fig 5). Similarly, the upregulation of the pro- inflammatory *Illb* gene at 4 h postmortem was followed by: upregulation of pro-inflammatory *Tnfrsf19, Tnf, Gpr31* and *Il8* genes and anti-inflammatory *Hmoxl* gene at 9 h, the upregulation of pro-inflammatory *Tnfa, Nfkbiaa*, and *Mknk2b* genes at 12 h, and the upregulation of anti-inflammatory *Crhr1* gene at 24 h. Of note, while none of the pro-inflammatory genes were upregulated past 48 h, the anti-inflammatory *Crhrl* gene remained upregulated at 48 h. It should also be noted that the *Il1b, Il8*, and *Tnfa* genes have been reported to be upregulated in traumatic impact injuries in postmortem tissues from human brains [87].

In the mouse, upregulated inflammation genes included: mitogen-activated protein kinase (*Map3k2*), TNF receptors (*Tnfrsf9, Tnfrs14*), B-cell lymphoma 6 protein (*Bcl6*), C-C chemokine receptor type 4 (Ccr4), Prokineticin-2 (*Prok2*), and platelet-activating factor receptor (*Pafr*) (Fig 5). The *Map3k2* gene encodes a kinase that activates pro-inflammatory NF-kB genes [85]. The *Tnfrsf9* and *Tnfrs14* genes encode receptor proteins that have pro-inflammatory functions [79]. The *Bcl6* gene encodes a transcription factor that has anti-inflammatory functions [88]. The *Ccr4* gene encodes a cytokine receptor protein associated with inflammation [70]. The *Prok2 gene* encodes a cytokine-like molecule, while the *Pafr* gene encodes a lipid mediator; both have pro-inflammatory functions [89,90].

Most inflammation-associated genes were upregulated within 1 h postmortem and continued to be upregulated for 12 to 48 h. The anti-inflammatory *Bcl6* gene was upregulated at two different times: 0.5 to 6 h and at 24 h suggesting that it is presumably being regulated by a feedback loop. It should also be noted that pro-inflammatory *Map3k2* and *Tnfrs14* genes were not upregulated after 24 and 12 h, respectively, which also suggests regulation by a putative feedback loop from the *Bcl6* gene product.

#### Summary of inflammation response

In both organisms, some of the upregulated genes have pro-inflammatory functions while others have anti-inflammatory functions, presumably reflecting regulation by feedback loops. The putative feedback loops involve an initial inflammatory reaction followed by an anti-inflammatory reaction to repress it [91]. The variation in the upregulation of these inflammatory genes suggests an underlying regulatory network is involved in organismal death.

### Apoptosis and related genes

Since apoptotic processes kill damaged cells for the benefit of the organism as a whole, we anticipated the upregulation of apoptosis genes in organismal death.

In the zebrafish, upregulated apoptosis genes included: Jun (*Jdp2, Jun*), Alkaline ceramidase 3 (*Acer3), Fos (Fosb, Fosab, Fosll*), IAP-binding mitochondrial protein A (*Diabloa*), Peroxiredoxin-2 (*Prdx2*), Potassium voltage-gated channel member 1 (*Kcnbl*), Caspase apoptosis-related cysteine peptidase 3b (*Casp3b*), DNA-damage-inducible transcript 3 (*Ddit3*), BCL2 (B-cell lymphomas 2)-interacting killer (*Bik*), and Ras association domain family 6 (*Rassf6*) (Fig 6). The *Jdp2* gene encodes a protein that represses the activity of the transcription factor activator protein 1 (*AP-1*) [92]. The *Acer3* gene encodes an enzyme that maintains cell membrane integrity/function and promotes apoptosis [93]. The *Fos* genes encode proteins that dimerize with *Jun* proteins to form part of the *AP-1* that promotes apoptosis [94,95]. The *Diabloa* gene encodes a protein that neutralizes inhibitors of apoptosis (IAP)-binding protein [95] and activates caspases [96]. The *Prdx2* gene encodes antioxidant enzymes that control cytokine-induced peroxide levels and inhibit apoptosis [97]. Although the *Kcnb1* gene encodes a protein used to make ion channels, the accumulation of these proteins in the membrane promotes apoptosis via cell signaling pathway [98]. The *Casp3b* encodes a protein that plays a role in the execution phase of apoptosis [99]. The *Ddit3* gene encodes a transcription factor that promotes apoptosis. The *Bik* gene encodes a protein that promotes apoptosis [100]. The *Rassf6* gene encodes a protein that promotes apoptosis [101].

**Fig 6.**
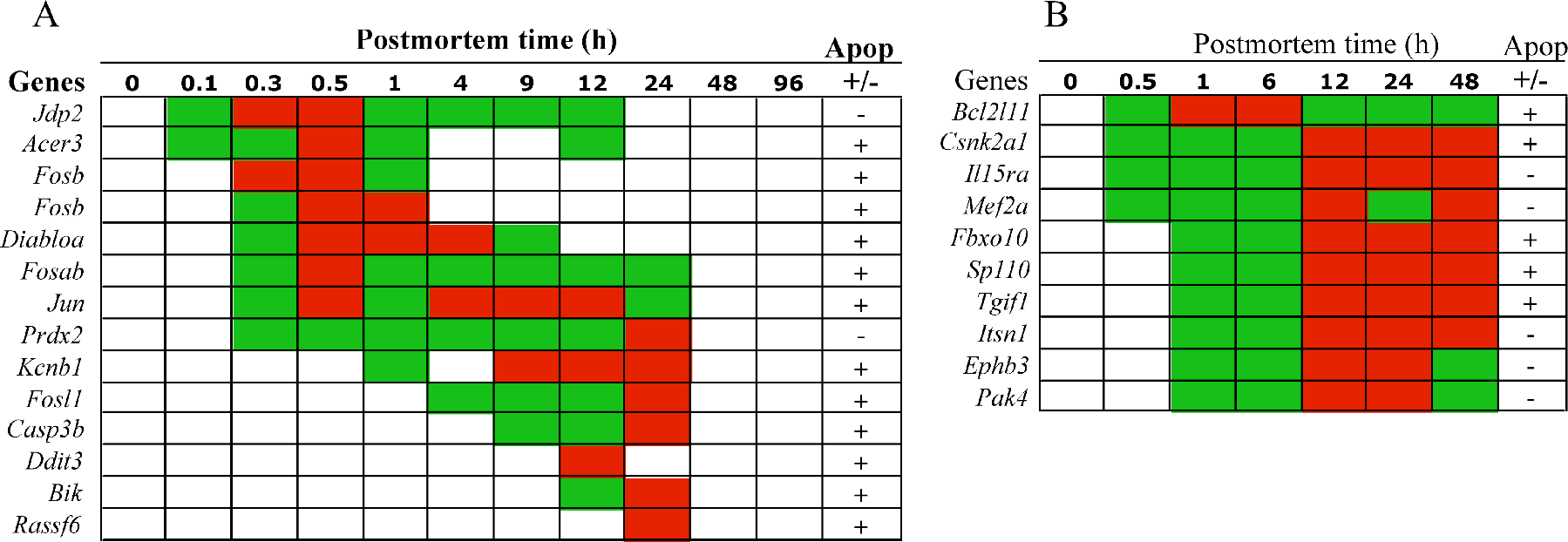
**Upregulated apoptosis genes by postmortem time (h). A, zebrafish; B, mouse. Apopotosis, Pro, +; Anti, − . Green, intermediate value; Red, maximum value. The *Fosb* gene was represented by two different probes**.

In the zebrafish, both anti-apoptosis *Jdp2* and pro-apopotosis *Acer3* genes were upregulated within 0.1 h postmortem (Fig 6). The upregulation of these genes was followed by the upregulation of five pro-apoptosis genes and one anti-apoptosis gene within 0.3 to 0.5 h. The transcriptional dynamics varied among the genes. Specifically, (i) the *Fosb* transcript was at low abundance after 1 h, (ii) the *Diabloa* and *Fosab* transcripts reached abundance maxima at 0.5 to 4 h and then were at low abundance after 9 h for the *Diabloa* and after 24 h for the *Fosab*, (iii) the *Jun* transcript reached two maxima (one at 0.5 and another at 4 to 12 h) - and after 24 h was at low abundance, (iv) the *Prdx2* transcript showed a continuous increase in abundance, reaching a maximum at 24 h and then was at low abundance. The remaining genes were pro-apoptosis and upregulated after 1 to 24 h postmortem. The *Ddit3* and *Rassf6* genes were very different from the other genes in that they were upregulated at one sampling time (12 h and 24 h, respectively) and then the transcripts were at low abundance. Apparently the apoptosis transcripts were at low abundance after 24 h in contrast to the transcripts in other categories (e.g. stress and immunity genes were upregulated for 96 h postmortem).

In the mouse, upregulated apoptosis-associated genes included: BCL2-like protein 11 (*Bcl2L11*), Casein kinase IIa (*Csnk2a1*), Interleukin 15 receptor subunit a (*Il15ra*), Myocyte enhancer factor 2 (Mef2a), F-box only protein 10 (*Fbxo10*), Sp110 nuclear body protein (*Sp110*), TGFB-induced factor homeobox 1 (*Tgif1*), Intersectin 1 (*Itsm1*) gene, the Ephrin type-B receptor 3 (*Ephb3*), and the p21 protein-activated kinase 4 (*Pak4*) (Fig 6) . The *Bcl2L11* gene encodes a protein that promotes apoptosis [102]. The *Csnk2a1* gene encodes an enzyme that phosphorylates substrates and promotes apoptosis [103]. The *Il15ra* gene encodes an anti-apoptotic protein [104]. The *Mef2a* gene encodes a transcription factor that prevents apoptosis [105]. The *Fbxo10* gene encodes a protein that promotes apoptosis [106]. The *Sp110* gene encodes a regulator protein that promotes apoptosis [107]. The *Tgif1* gene encodes a transcription factor that blocks signals of the transforming growth factor beta (*TGFfi*) pathway; and therefore, is pro-apoptosis [108]. The *Itsn1* gene encodes an adaptor protein that is anti-apoptosis [109]. The *Ephb3* gene encodes a protein that binds ligands on adjacent cells for cell signaling and suppresses apoptosis [104]. The *Pak4* gene encodes a protein that delays the onset of apoptosis [110].

In the mouse, pro- and anti-apoptosis genes were upregulated within 0.5 h postmortem - however, with exception to *Bcl2L11*, most reached transcript abundance maxima at 12 to 48 h postmortem (Fig 6). The *Bcl2L11* transcripts reached abundance maxima at 1 and 6 h postmortem.

#### Summary of apoptotic response

In both organisms, pro- and anti-apoptosis genes were upregulated in organismal death. However, the timings of the upregulation, the transcript abundance maximum, and the duration of the upregulation varied by organism. The results suggest the apoptotic genes and their regulation are distinctly different in the zebrafish and the mouse, with the mouse genes being upregulated up to 48 h postmortem, while zebrafish genes are upregulated for 24 h. Nonetheless, the pro- and anti-apoptosis genes appear to be inter-regulating each another.

### Transport gene response

Transport processes maintain ion/solute/protein homeostasis and are involved in influx/efflux of carbohydrates, proteins, signaling molecules, and nucleic acids across membranes. We anticipated that transport genes should be upregulated in organismal death in response to dysbiosis.

In the zebrafish, upregulated transport-associated genes included: Solute carrier family 26 Anion Exchanger Member 4 (*Slc26a4*), Potassium channel voltage gated subfamily H (*Kcnh2*), Transmembrane Emp24 domain containing protein 10 (*Tmed10*), Leucine rich repeat containing 59 (*Lrrc59*), the Nucleoprotein TPR (*Tpr*), Importin subunit beta-1 (*Kpnb1*), Transportin 1 (*Tnpo1*), Syntaxin 10 (*Stx10*) and Urea transporter 2 (*Slc14a2*) (Fig 7). Of note, the four *Tmed10* transcripts shown in Fig 7, each represents a profile targeted by an independent probe. The transcription profiles of this gene were identical, indicating the high reproducibility of the Gene Meter approach. The *Slc26a4* gene encodes prendrin that transports negatively-charged ions (i.e., Cl^-^, bicarbonate) across cellular membranes [111]. The *Kcnh2* gene encodes a protein used to make potassium channels and is involved in signaling [112]. The *Tmed10* gene encodes a membrane protein involved in vesicular protein trafficking [113]. The *Lrrc59, Tpr, Tnpol*, and *Kpnbl* genes encode proteins involved in trafficking across nuclear pores [114,115,116,117]. The *Stx10* gene encodes a protein that facilitates vesicle fusion and intracellular trafficking of proteins to other cellular components [118]. The *Slc14a2* gene encodes a protein that transports urea out of the cell [119].

**Fig 7.**
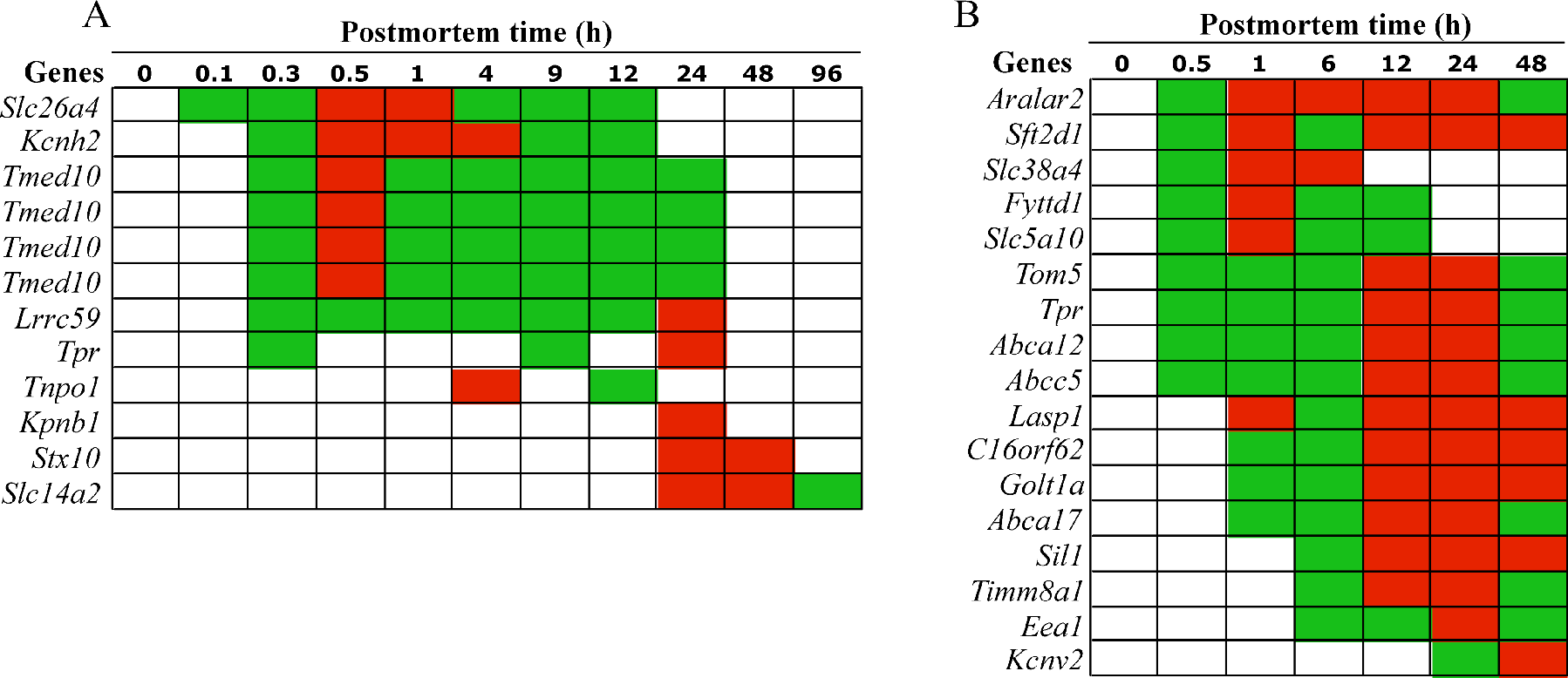
**Upregulated transport genes by postmortem time (h) and stress category. A. zebrafish; B, mouse. Green, intermediate value; Red, maximum value. The *Tmed10* gene was represented by four different probes**.

The *Slc26a4, Kcnh2, Lrrc59* and *Tpr* genes were initially upregulated within 0.3 h postmortem and continue to be expressed for 12 to 24 h. The *Tnpo1* gene was upregulated at two times: 4 h and 12 h suggesting putative regulation by a feedback loop. The remaining genes were upregulated at 24 h. The upregulation of the *Slc14a2* gene suggests a build up of urea in zebrafish cells at 24 to 96 h postmortem, which could be due to the accumulation of urea under hypoxic conditions by the *Arg2* gene (see *Hsp* stress response section).

In the mouse, the following transport-associated genes were upregulated: Calcium-bind mitochondrial carrier protein (*Aralar2*), Sodium-coupled neutral amino acid transporter 4 (*Slc38a4*), SFT2 domain containing 1 (*Sft2d1*), Uap56-interacting factor (*Fyttd1*), Solute carrier family 5 (sodium/glucose co-transporter) member 10 (*Slc5a10*), Mitochondrial import receptor subunit (*Tom5*), ‘Translocated promoter region’ (*Tpr*), ATP-binding cassette transporter 12 (*Abca12*), Multidrug resistant protein 5 (Abc5), LIM and SH3 domain-containing protein (*Lasp1*), Chromosome 16 open reading frame 62 (*C16orf62*), Golgi transport 1 homolog A (*Golt1a*), ATP-binding cassette transporter transporter 17 (*Abca17*), Nucleotide exchange factor (*Sil1*), Translocase of inner mitochondrial membrane 8A1 (*Timm8a1*), Early endosome antigen 1 (*Eea1*), and Potassium voltage-gated channel subfamily V member2 (*Kcnv2*) (Fig 7). The *Aralar2* gene encodes a protein that catalyzes calcium-dependent exchange of cytoplasmic glutamate with mitochondrial aspartate across the mitochondrial membrane and may function in the urea cycle [120]. The *Slc38a4* gene encodes a symport that mediates transport of neutral amino acids and sodium ions [121]. The *Sft2d1* gene encodes a protein involved in transporting vesicles from the endocytic compartment of the Golgi complex [122]. The *Fyttd1* gene is responsible for mRNA export from the nucleus to the cytoplasm [123]. The *Slc5a10* gene encodes a protein that catalyzes carbohydrate transport across cellular membranes [124]. The *Tom5* gene encodes a protein that plays a role in importation to proteins destined to mitochondrial sub-compartments [125]. The *Abca12*, *Abca17* and *Abc5* genes encode proteins that transport molecules across membranes [126,127,128]. The *Lasp1* gene encodes a protein that regulates ion-transport [129]. The *C16orf62* gene encodes a protein involved in protein transport from the Golgi apparatus to the cytoplasm [130]. The *Golt1a* gene encodes a vesicle transport protein [122]. The *Sil1* gene encodes a protein involved in protein translocation into the endoplasmic reticulum [131]. The *Timm8a1* gene encodes a protein that assists importation of other proteins across inner mitochondrial membranes [132]. The *Eea1* gene encodes a protein that acts as a tethering molecule for vesicular transport from the plasma membrane to the early endosomes [133]. The *Kcnv2* gene encodes a membrane protein involved in generating action potentials [134].

Within 0.5 h postmortem, genes involved in: (i) ion and urea regulation (*Aralar*), (ii) amino acid (*Slc38a4*), carbohydrate (*Slc5a10*), and protein (*Sft2d1, Tom5*) transport, (iii) mRNA nuclear export (*Fyttd1, Tpr*), and (iv) molecular efflux (*Abca12, Abc5*) were upregulated in the mouse. The transcription profiles of these genes varied in terms of transcript abundance maxima and duration of the upregulation. While the transcripts of *Aralar, Sft2d1, Slc38a4, Fyttd1* and *Slc5a10* reached abundance maxima at 1 h, those of *Tom5, Tpr, Abca12*, and *Abc5* reached maxima at 12 to 24 h postmortem. The duration of the upregulation also varied for these genes since most were upregulated for 48 h postmortem, while the *Sft2d1, Fyttd1* and *Slc5a10* were upregulated from 0.5 to 12 h or more. The shorter duration of upregulation suggests prompt gene repression. The transcript abundance of *Lasp1, C16orf62, Golt1a*, and *Abca17* increased at 1 h postmortem and remained elevated for 48 h. The transcripts of *Sil1, Timm8a1, Eea1* increased in abundance at 6 h, while those of *Kcnv2* increased at 24 h postmortem and remained elevated for 48 h.

#### Summary of transport genes

The upregulation of transport genes suggests attempts by zebrafish and mice to reestablish homeostasis. Although half of these genes were upregulated within 0.5 h postmortem, many were upregulated at different times and for varying durations. While most of the transport gene transcripts in the zebrafish were at low abundance after 24 h, most transport gene transcripts in the mouse remained at high abundance at 24 to 48 h postmortem.

### Development genes

An unexpected finding in this study was the discovery of upregulated development genes in organismal death. Most development genes are involved at the early stage of development in the zebrafish and mouse; therefore, we did not anticipate their upregulation in organismal death.

In the zebrafish, upregulated development genes included: LIM domain containing protein 2 (*Limd2*), Disheveled-associated activator of morphogenesis 1 (*Daam1b*), Meltrin alpha (*Adam12*), Hatching enzyme 1a (*He1a*), Midnolin (*Midn*), Immediate early response 2 (*Ier2*), Claudin b (*Cldnb*), Regulator of G-protein signaling 4-like (*Rgs4*), Proline-rich transmembrane protein 4 (*Prrt4*), Inhibin (*Inhbaa*), Wnt inhibitory factor 1 precursor (*Wif1*), Opioid growth factor receptor (*Ogfr*), Strawberry notch homolog 2 (*Sbno2*), and Developing brain homeobox 2 (*Dbx2*) (Fig 8). The *Limd2* gene encodes a binding protein that plays a role in zebrafish embryogenesis [135]. The *Daam1b* gene regulates endocytosis during notochord development [136]. The *Adam12* gene encodes a metalloprotease-disintegrin involved in myogenesis [137]. The *He1a* gene encodes a protein involved in egg envelope digestion [138]. The*Midn* gene encodes a nucleolar protein expressed in the brain and is involved in the regulation of neurogenesis [139,140]. The *Ier2* gene encodes a protein involved in left-right asymmetry patterning in the zebrafish embryos [141]. The *Cldnb* gene encodes a tight junction protein in larval zebrafish [142]. The *Rgs4* gene encodes a protein involved in brain development [143]. The *Prrt4* gene encodes a protein that is predominantly expressed in the brain and spinal cord in embryonic and postnatal stages of development. The *Inhbaa* gene encodes a protein that plays a role in oocyte maturation [144]. The *Wif1* gene encodes a WNT inhibitory factor that controls embryonic development [145]. The *Ogfr* gene plays a role in embryonic development [146]. The *Sbno2* gene plays a role in zebrafish embryogenesis [147]. The *Dbx2* gene encodes a transcription factor that plays a role in spinal cord development [148].

**Fig 8.**
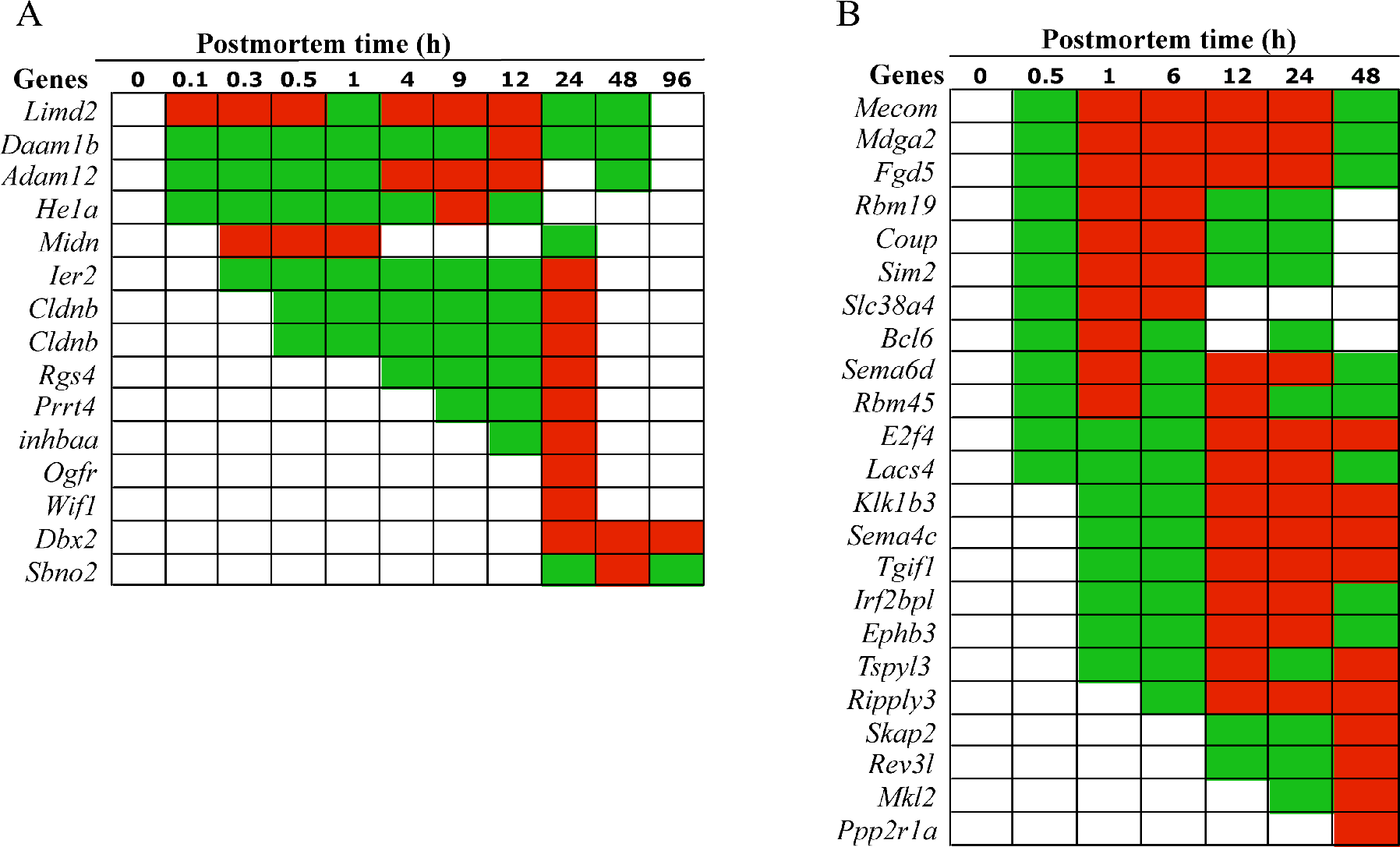
**Upregulated development genes by postmortem time (h) and stress category. A. zebrafish; B, mouse. Green, intermediate value; Red, maximum value. The *Cldnb* gene was represented by two different probes**.

Although the abundances of *Limd2, Daaml, Adam12, and Hela* transcripts increased in the zebrafish within 0.1 h postmortem, other gene transcripts in this category increased from 0.3 to 24 h postmortem reaching abundance maxima at 24 h or more.

In the mouse, development genes were upregulated included: MDS1 and EVI1 complex locus protein EVI1 (*Mecom*), MAM domain containing glycosylphosphatidylinositol anchor 2 (*Mdga2*), FYVE, RhoGEF and PH domain containing 5 (*Fgd5*), RNA binding motif protein 19 (*Rbm19*), Chicken ovalbumin upstream promoter (*Coup*), Single minded homolog 2 (*Sim2*), Solute carrier family 38, member 4 (*Slc38a4*), B-cell lymphoma 6 protein (*Bcl6*), Sema domain transmembrane domain (TM) cytoplasmic domain (semaphorin) 6D (*Sema6d*), RNA binding motif protein *45(Rbm45*), Transcription factor E2F4 (*E2f4*), Long chain fatty acid- CoA Ligase 4 (*Lacs4*), Kallikrein 1-related peptidase b3 (*Klk1b3*), Sema domain, Immunoglobulin domain, transmembrane domain and short cytoplasmic domain (*Sema4c*), TGFB-induced factor homeobox 1 (*Tgif1*), Interferon regulatory factor 2-binding protein-like (*Irf2bpl*), Ephrin type-B receptor 3 (*Ephb3*), Testis-specific Y-encoded-like protein 3 (*Tspyl3*), Protein ripply *3(Ripply3*), Src kinase-associated phosphoprotein 2 (Skap2), DNA polymerase zeta catalytic subunit (*Rev3l*), MKL/Myocardin-Like 2 (*Mkl2*), and Protein phosphatase 2 regulatory subunit A (*Ppp2r1a*) (Fig 8). The *Mecom* gene plays a role in embryogenesis and development [149]. The *Mdga2* gene encodes immunoglobins involved in neural development [150]. The *Fgd5* gene is needed for embryonic development since it interacts with hematopoietic stem cells [151]. The *Rbm19* gene is essential for preimplantation development [152]. The *Coup* gene encodes a transcription factor that regulates development of the eye [153] and other tissues [154]. The *Sim2* gene encodes a transcription factor that regulates cell fate during midline development [155]. The *Slc38a4* gene encodes a regulator of protein synthesis during liver development and plays a crucial role in fetal growth and development [156,157]. The *Bcl6* gene encodes a transcription factor that controls neurogenesis [158]. The *Sema6d* gene encodes a protein involved in retinal development [159]. The *Rbm45* gene encodes a protein that has preferential binding to poly(C) RNA and is expressed during brain development [160]. The *E2f4* gene is involved in maturation of cells in tissues [161]. The *Lacs4* gene plays a role in patterning in embryos [162]. The *Klk1b3* gene encodes a protein that plays a role in developing embryos [163]. The *Sema4c* gene encodes a protein that has diverse functions in neuronal development and heart morphogenesis [164, 165]. The *Tgif1* gene encodes a transcription factor that plays a role in trophoblast differentiation [166]. The *Irf2bpl* gene encodes a transcriptional regulator that plays a role in female neuroendocrine reproduction [167]. The *Ephb3* gene encodes a kinase that plays a role in neural development [168]. The *Tspyl3* gene plays a role in testis development [169]. The *Ripply3* gene encodes a transcription factor involved in development of the ectoderm [170]. The *Skap2* gene encodes a protein involved in actin reorganization in lens development [171]. The *Rev3l*gene encodes a polymerase that can replicate past certain types of DNA lesions and is necessary for embryonic development [172]. The*Mkl2* gene encodes a transcriptional co-activator is involved in the formation of muscular tissue during embryonic development [173]. The *Ppp2r1a* gene plays a role in embryonic epidermal development [174].

While the *Mecom, Mdga2, Fgd5, Rbm19, Coup, Sim2, Slc38a4, Bcl6, Sema6d, Rbm45, E2f4* and *Lacs4* genes were upregulated within 0.5 h postmortem, the other genes were upregulated at 1 h to 48 h with most transcripts reaching abundance maxima at 12 h or more.

#### Summary of development genes

In organismal death, there is progressive activation of developmental genes suggesting that they are no longer silenced. A possible reason for this activation is that postmortem physiological conditions resemble those during development.

### Cancer genes

There are a number of databases devoted to cancer and cancer-related. Upon cross-referencing the genes found in this study, we discovered a significant overlap. The genes found in this search are presented below.

In the zebrafish, the following cancer genes were upregulated: *Jdp2*, xanthine dehydrogenase (*Xdh*), *Egr1, Adam12*, myosin-IIIa (*Myo3a*), *Fosb, Jun*, Integrin alpha 6b (*Itga6), Ier2, Tpr*, Dual specificity protein phosphatase 2 (*Dusp2*), Disintegrin and metallopeptidase domain 28 (*Adam28), Tnpo1*, Ral guanine nucleotide dissociation stimulator-like (*Rgl1*), Carcinoembryonic antigen-related cell adhesion molecule 5 (*Ceacaml), Fosl1, Il1b, Hif1a*, Serine/threonine-protein phosphatase 2A regulatory (*Ppp2r5d*), DNA replication licensing factor (*Mcm5), Gadd45*, Myosin-9 (*Myh9), Casp3, Tnf, Il8*, Cyclic AMP-dependent transcription factor (*Atf3*), small GTPase (RhoA), *Mknk2*, Ephrin type-A receptor 7 precursor (*Epha7*), ETS-related transcription factor (*Elf3), Nfkbia, Kpnb1, Wif1*, RAS guanyl-releasing protein 1 (*Rasgrp*), Ras association domain-containing protein 6 (*Rassf6), Cyba*, DNA-damage-inducible transcript 3 (*Ddit3*), Serine/threonine-protein kinase (*Sbk1*), and Tyrosine-protein kinase transmembrane receptor (*Ror1*) (Fig 9).

**Fig 9.**
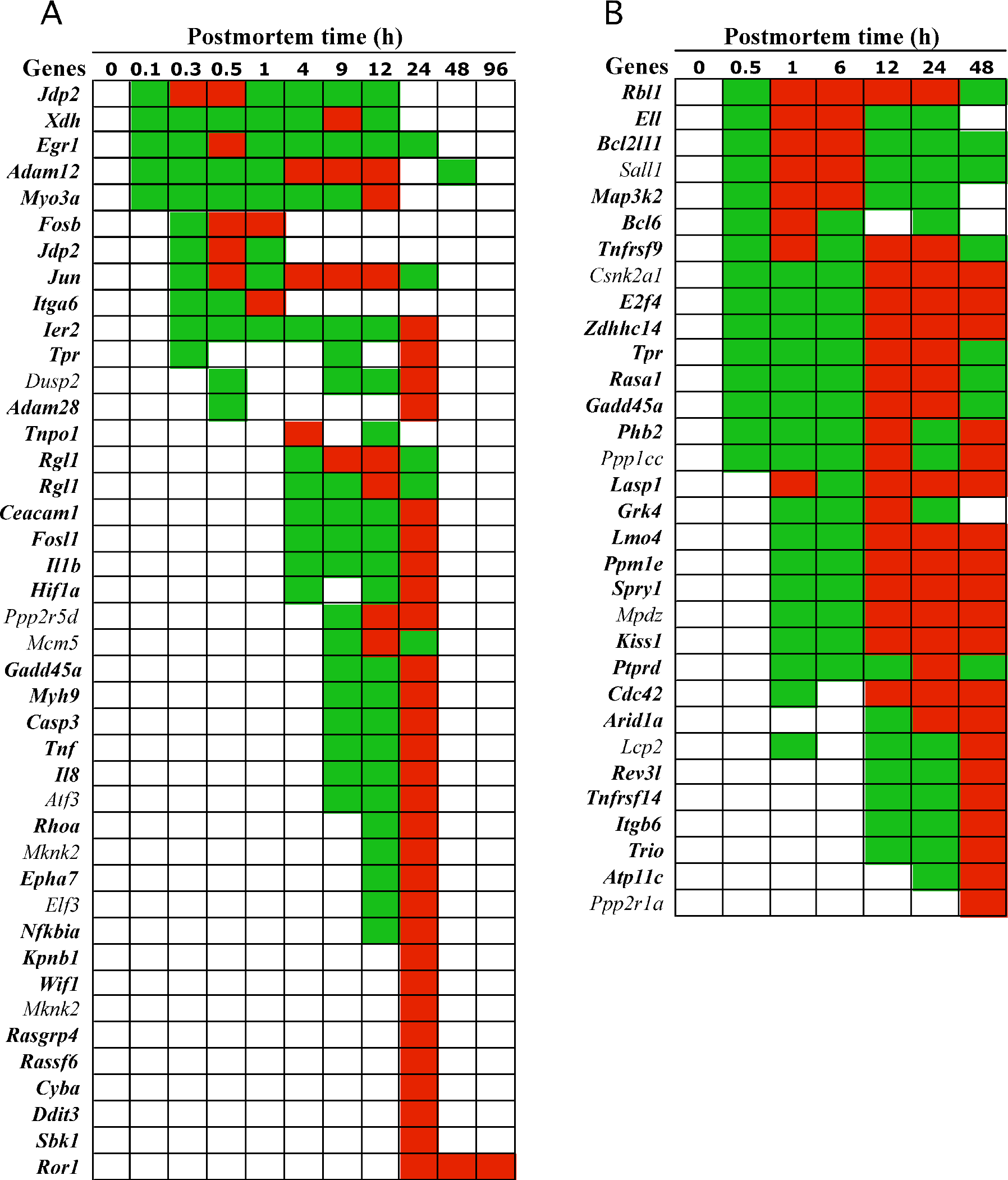
**Upregulated cancer genes by postmortem time (h). A, Zebrafish; B, Mouse. Green, intermediate value; Red, maximum value. Bold gene name means it was found in more than one cancer database. The *Rgll* gene was represented by two different probes**.

In the mouse, the following cancer genes were upregulated: retinoblastoma-like protein 1 (*Rbl1*), Elongation factor RNA polymerase II (*Ell*), Bcl-2-like protein 11 (*Bcl2l11*), Sal-like protein 1 (*Sall1), Map3k2, Bcl6, Tnfrsf9*, CK2 target protein 2 (*Csnk2a1*), Transcription factor E2f4 (*E2f4*), Zinc finger DHHC-type containing 14 (*Zdhhc14), Tpr*, RAS p21 protein activator 1 (*Rasa1), Gadd45*, prohibitin (*Phb2*), Serine/threonine-protein phosphatase PP1-gamma catalytic (Ppp1cc), *Lasp1*, G protein-coupled receptor kinase 4 (*Grk4*), LIM domain transcription factor (*Lmo4*), Protein phosphatase 1E (*Ppm1e*), Protein sprouty homolog 1 (*Spry1*), Multiple PDZ domain protein (*Mpdz*), Kisspeptin receptor (*Kiss1*), Receptor-type tyrosine-protein phosphatase delta precursor (*Ptprd*), Small effector protein 2-like (*Cdc42*), AT-rich interactive domain-containing protein 1A (*Arid1a*), Lymphocyte cytosolic protein 2 (*Lcp2*), DNA polymerase zeta catalytic subunit (*Rev3l), Tnfrsf14*, Integrin beta-6 precursor (*Itgb6*), Triple functional domain protein (*Trio*), ATPase class VI type 11C (*Atp11c*), and Serine/threonine-protein phosphatase 2A regulatory (*Ppp2r1a*) (Fig 9).

#### Summary of cancer genes

These genes were classified as “cancer genes” in a Cancer Gene Database [7] (Fig 9). The timing, duration and peak transcript abundances differed within and between organisms. Note that some transcripts had two abundance maxima. In the zebrafish, this phenomenon occurred for *Adam12, Jun, Tpr, Dusp2, Tnpo1*, and *Hif1a* and in the mouse, *Bcl6, Tnfrs9, Lasp1, Cdc42*, and *Lcp2*, and is consistent with the notion that the genes are being regulated through feedback loops.

### Epigenetic regulatory genes

Epigenetic regulation of gene expression involves DNA methylation and histone modifications of chromatin into active and silenced states [175]. These modifications alter the condensation of the chromatin and affect the accessibility of the DNA to the transcriptional machinery. Although epigenetic regulation plays important role in development, modifications can arise stochastically with age or in response to environmental stimuli [176]. Hence, we anticipated that epigenetic regulatory genes should be involved in organismal death.

In the zebrafish, the following epigenetic genes were upregulated: Jun dimerization protein 2 (*Jdp2*), Chromatin helicase protein 3 (*Chd3*), Glutamate-rich WD repeat-containing protein 1 (*Grwd1*), Histone H1 (*Histh1l*), Histone cluster 1, H4-like (*Hist1h46l3*) and Chromobox homolog 7a (*Cbx7a*) (Fig 10). The *Jdp2* gene is thought to inhibit the acetylation of histones and repress expression of the c-Jun gene [177]. The *Chd3* gene encodes a component of a histone deacetylase complex that participates in the remodeling of chromatin [178]. The *Grwd1* gene is thought to be a histone-binding protein that regulates chromatin dynamics at the replication origin [179]. The *Histh1l* gene encodes a histone protein that binds the nucleosome at the entry and exit sites of the DNA and the *Hist1h46l3* gene encodes a histone protein that is part of the nucleosome core [180]. The *Cbx7a* gene encodes an epigenetic regulator protein that binds noncoding RNA and histones and represses gene expression of a tumor suppressor [181].

**Fig 10.**
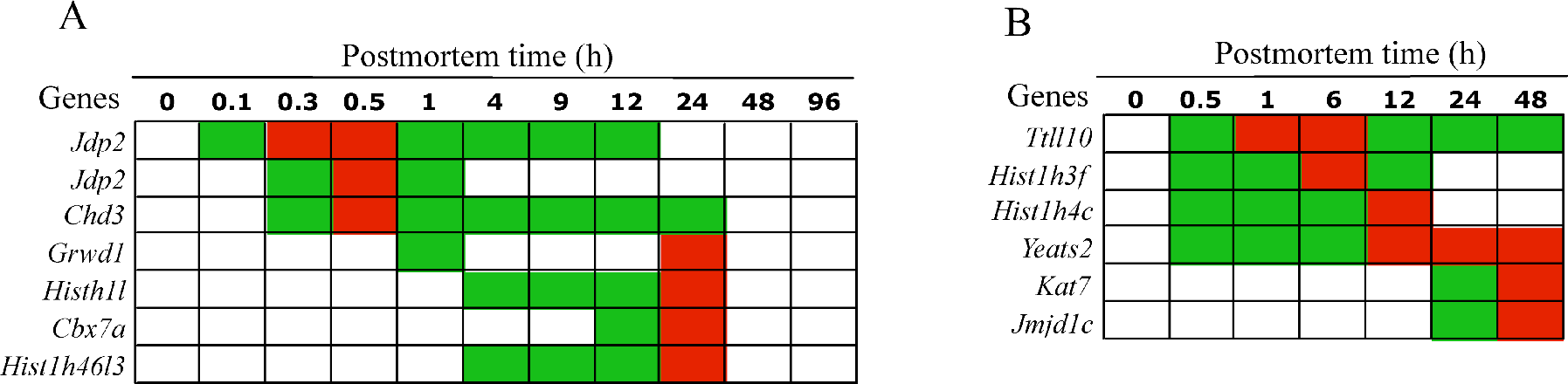
**Upregulated epigenetic genes by postmortem time (h). A, Zebrafish; B, Mouse. Green, intermediate value; Red, maximum value. Bold gene name means it was found in more than one cancer database. The *Jdp2* gene was represented by two different probes**.

Both *Jdp2* and *Chd3* genes were upregulated within 0.3 h postmortem, with their transcripts reaching abundance maxima at 0.5 h. Note that two different probes targeted the *Jdp2* transcript. The *Grwdl* gene was upregulated at 1 h and 24 h postmortem. The histone genes were upregulated at 4 h postmortem with their transcripts reaching abundance maxima at 24 h. The *Cbx7a* gene was upregulated at 12 h and its transcript reached an abundance maximum at 24 h. Transcripts of these genes were at low abundance after 24 h.

In the mouse, the following epigenetic genes were upregulated: Tubulin tyrosine ligase-like family member 10 (*TtlllO*), Histone cluster 1 H3f (*Hist1h3f*), Histone cluster 1 H4c (*Hist1h4c*), YEATS domain containing 2 (*Yeats2*), Histone acetyltransferase (*Kat7*), and Probable JmjC domain-containing histone demethylation protein 2C (*Jmjd1c*) (Fig 10). The *Ttll10* gene encodes a polyglycylase involved in modifying nucleosome assembly protein 1 that affects transcriptional activity, histone replacement, and chromatin remodeling [182]. The *Hist1h3f* and *Hist1h4c* genes encode histone proteins are the core of the nucleosomes [183]. The *Yeats2* gene encodes a protein that recognizes histone acetylations so that it can regulate gene expression in the chromatin [184]. The *Kat7* gene encodes an acetyltransferase that is a component of histone binding origin-of-replication complex, which acetylates chromatin and therefore regulates DNA replication and gene expression [185]. The *Jmjd1c* gene encodes an enzyme that specifically demethylates ‘Lys-9′ of histone H3 and is implicated in the reactivation of silenced genes [186].

The *Ttll10*, *Yeats2* and histone protein genes were upregulated 0.5 h postmortem but their transcripts reached abundance maxima at different times with the *Ttll10* transcript reaching a maximum at 1 to 6 h, the histone transcripts reaching maxima at 6 and 12 h postmortem, and the *Yeats2* transcript reaching maxima at 12 to 24 h postmortem (Fig 10). The *Kat7* and *Jmjd1c* genes were upregulated at 24 h and their transcripts reached abundance maxima at 48 h postmortem.

#### Summary of epigenetic regulatory genes

The upregulation of genes encoding histone proteins, histone-chromatin modifying proteins, and proteins involved in regulating DNA replication at the origin were common to the zebrafish and the mouse. These findings indicate that epigenetic regulatory genes are modifying chromatin structure by regulating the accessibility of transcription factors to the promoter or enhancer regions in organismal death.

### Percentage of upregulated genes with postmortem time

The % of upregulated genes was defined as the number of upregulated genes at a specific postmortem time over the total number of upregulated genes in a category. A comparison of the % of upregulated genes by postmortem time of all upregulated genes revealed similarities between the zebrafish and the mouse. Specifically, most genes were upregulated between 0.5 to 24 h postmortem, and after 24 h, the upregulation of most genes stopped (Fig 11, “All genes”). It should be noted that the same pattern was found in stress, transport and development categories for both organisms. However, in the zebrafish, the immunity, inflammation, apoptosis and cancer categories differed from the mouse. Specifically, the genes in the immunity, inflammation, and cancer categories were upregulated much later (1 to 4 h) in the zebrafish than the mouse, and the duration of upregulation was much shorter. For example, while 90% of the genes in the immunity and inflammation categories were upregulated in the mouse within 1 h postmortem, less than 30% of the genes were upregulated in the zebrafish (Fig 11), indicating a slower initial response. It should be noted that while the number of upregulated immunity genes reaching transcript abundance maxima occurred at 24 h postmortem in both organisms, the number of inflammation genes reaching transcript abundance maxima occurred at 1 to 4 h in the mouse, and 24 h in the zebrafish. The significance of these results is that the inflammation response occurs rapidly and robustly in the mouse while in the zebrafish, it takes longer to establish, which could be attributed to phylogenetic differences. There were significant differences in the upregulation of apoptosis genes between the zebrafish and the mouse. In the mouse, the number of upregulated apoptosis genes reached 100% at 1 h postmortem and remained sustained for 48 h postmortem while the % of upregulated genes in the zebrafish never reached 70% and gene upregulation was abruptly stopped after 12 h.

**Fig 11.**
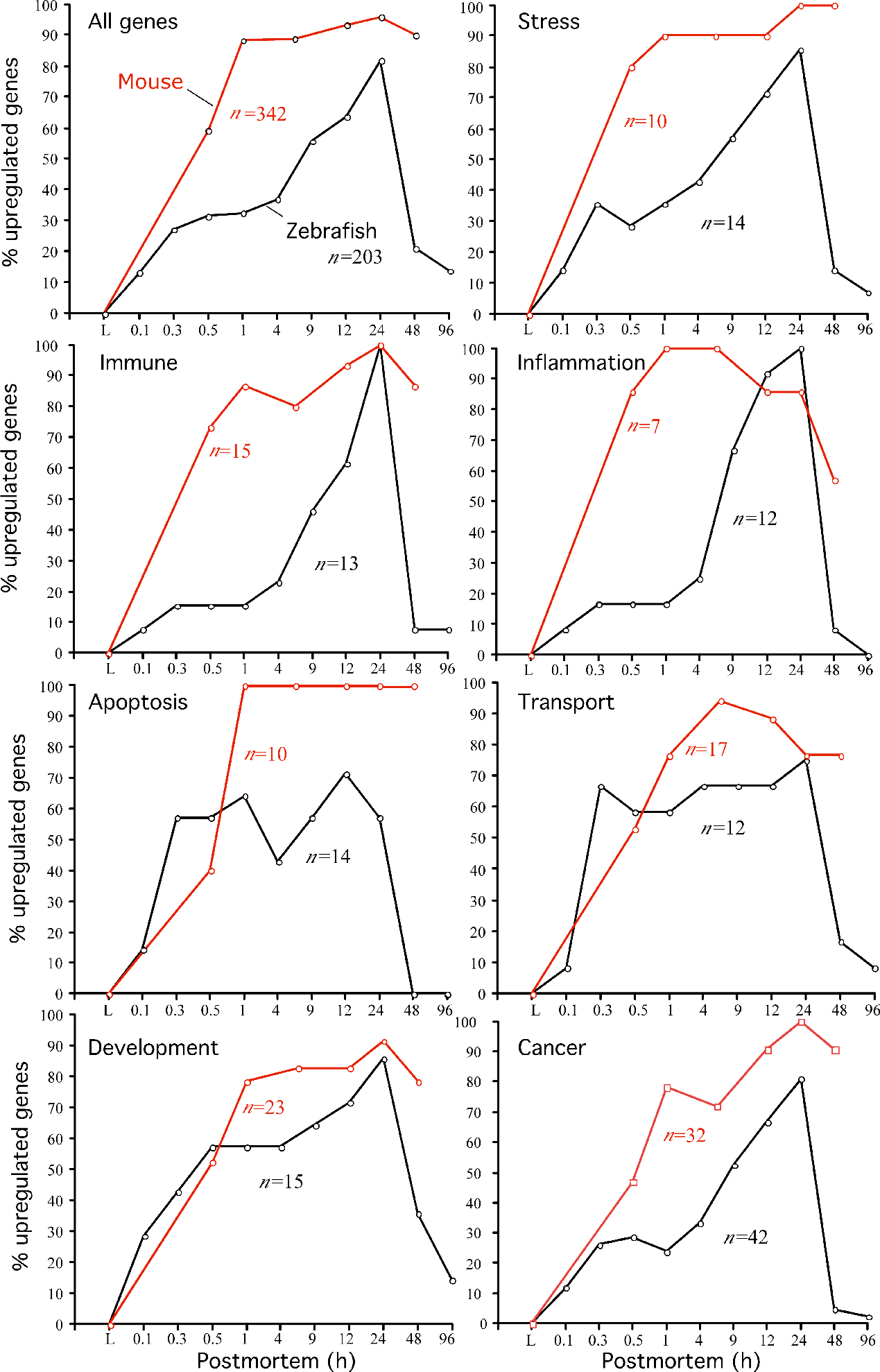
**% of upregulated genes by postmortem time and category. Number of total genes by organism and category are shown. “All genes” refers to the genes that significantly contributed to the ordination plots. Mouse is red; Zebrafish is black**.

## DISCUSSION

### Active gene expression or residual transcription levels?

One could argue that our study identifies only residual transcription levels of presynthesized mRNA (rather than newly-synthesized mRNA) in dead tissues that happen to be enriched with postmortem time. In other words, the observed upregulation of genes may be viewed as an artifact merely reflecting the “enrichment of specific mRNA transcripts” (e.g. stable mRNA) with time. The data, however, does not support this idea because if it were true, one would expect stable transcripts to monotonically increase with time, as they become more enriched (higher abundances) with postmortem time. The data show that the transcripts of most upregulated genes did not display monotonic behavior; rather, the transcripts reached abundance maximum (peak) or maxima (peaks) at various postmortem times (Fig 2D and 2E). This finding should not be a surprise because a statistical procedure was implemented to detect genes that were significantly upregulated - which is essentially selecting gene transcriptional profiles that had peaks. The statistics for the procedure was calibrated with more than a billion simulations. The simulation process corrected for multiple comparisons. The residual transcription level and enrichment idea is also not supported by transcriptional profiles displaying an up-, down-, and up- regulation pattern, which putatively indicates feedback loops ((Fig S3). Similarly, simple differential decay rates of mRNAs would not display this pattern because a transcript cannot be stable at one postmortem time, unstable at a subsequent time, and then stable again. A mRNA transcript is either stable or it is not. In conclusion, the gene profiles showing upregulation are the results of active transcription.

It should be noted that postmortem upregulation of genes has been previously reported in cadavers. Using reverse transcription real-time quantitative PCR (RT-RTqPCR), a study showed significant increases in expression of myosin light chain 3 (*Myl3*), matrix metalloprotease 9 (*Mmp9*), and vascular endothelial growth factor A (*Vegfa*) genes in body fluids after 12 h postmortem (187). Interestingly, postmortem upregulation of myosin-related and matrix metalloprotease genes was also found in our study. Specifically, the myosin-related genes included: Myosin-Ig (*Myo1g*) in the mouse, and Myosin-IIIa (*Myo3a*) and Myosin-9 (*Myh9*) in the zebrafish. The matrix metalloproteinase gene included the metalloproteinase-14 (*Mmp14b*) gene in the zebrafish. The *Myo1g* gene encodes a protein regulating immune response (189), the Myosin-IIIa (*Myo3a*) gene encodes an uncharacterized protein, the Myosin-9 (*Myh9*) gene encodes a protein involved in embryonic development (190), and the *Mmp14b* gene encodes an enzyme regulating cell migration during zebrafish gastrulation (188). The *Myo1g, Myh9* and *Mmp14b* transcripts increased right after death and reached abundance maxima at 24 h postmortem, while the *Myo3a* transcript reached an abundance maximum at 12 h postmortem. The significance of these results is two-fold: (i) two different technologies (RT-RTqPCR and Gene Meters) have now demonstrated active postmortem upregulation of genes and this expression has now been reported in three organisms (human, zebrafish, and mouse), and (ii) there might be significant overlap in genes upregulated in death as we have showed with myosin- and matrix metalloprotease genes, which warrants further studies using other vertebrates. The purpose of such studies would be to understand common mechanisms involved in the shutdown of highly ordered biological systems.

### Why study gene expression in death?

The primary motivation for the study was driven by curiosity in the processes of shutting down a complex biological system – which has received little attention so far. While the development of a complex biological system requires time and energy, its shutdown and subsequent disassembly entails the dissipation of energy and unraveling of the complex structures, which does not occur instantaneously and could provide insights into interesting paths. Moreover, other fields of research have examined the shutdown of complex systems (e.g., societies [191], government [192], electrical black outs [193]). Yet, to our knowledge, no study has examined long-term postmortem gene expression of vertebrates kept in their native conditions. The secondary motivation for the study was to demonstrate the precision of Gene Meter technology for gene expression studies to biologists who believe that high throughput DNA sequencing is the optimal approach.

### Thermodynamic sinks

We initially thought that sudden death of a vertebrate would be analogous to a car driving down a highway and running out of gas. For a short time, engine pistons will move up and down and spark plugs will spark -- but eventually the car will grind to a halt and “die”. Yet, in our study we find hundreds of genes are upregulated many hours postmortem, with some (e.g., *Kcnv2, Pafr, Degs2, Ogfod1, Ppp2rla, Ror1*, and *Iftm1*) upregulated days after organismal death. This finding is surprising because in our car analogy, one would not expect window wipers to suddenly turn on and the horn to honk several days after running out of gas.

Since the postmortem upregulation of genes occurred in both the zebrafish and the mouse in our study, it is reasonable to suggest that other multicellular eukaryotes will display a similar phenomenon. What does this phenomenon mean in the context of organismal life? We conjecture that the highly ordered structure of an organism – evolved and refined through natural selection and self-organizing processes – undergoes a thermodynamically driven process of spontaneous disintegration through complex pathways, which apparently involve the upregulation of genes and feedback loops. While evolution played a role in pre-patterning of these pathways, it does not play any role in its disintegration fate. One could argue that some of these pathways have evolved to favor healing or “resuscitation” after severe injury. For example, the upregulation of inflammation response genes indicate that a signal of infection or injury is sensed by the still alive cells after death of the body. Alternatively, the upregulation may be due to fast decay of some repressors of genes or whole pathways (see below). Hence, it will be of interest to study this in more detail, since this could, for example, provide insights into how to better preserve organs retrieved for transplantation.

### Chemical automator – on the way down to equilibrium

As one would expect, a living system is a collection of chemical reactions linked together by the chemicals participating in them. Having these reactions to depend on one another to a certain extent, we conjecture that the observed upregulation of genes is due to thermodynamic and kinetic controls that are encountered during organismal death. For example, epigenetic regulatory genes that were upregulated included histone modification genes (e.g., *Histh1l*) and genes interacting with chromatin (e.g., *Grwd1*, *Chd3*, *Yeats*, *Jmjd1c*) (Fig 10). It is possible that the activation of these genes was responsible for the unraveling of the nucleosomes, which enabled transcription factors and RNA polymerases to transcribe developmental genes that have been previously silenced since embryogenesis. The energy barrier in this example is the tightly wrapped nucleosomes that previously did not allow access to developmental genes. Other energy or entropy barriers could be nucleopores that allow the exchange of mRNA and other molecules between the mitochondria and the cytosol (e.g., *Tpr, Tnpo1, Lrrc59*), or the ion/solute protein channels (e.g., *Aralar2, Slc38a4*) that control intracellular ions that regulate apoptotic pathways [194,195].

The upregulation of genes indicates new molecules were synthesized. Hence, there was sufficient energy and resources (e.g., RNA polymerase, dNTPs) in dead organisms to maintain gene transcription to 96 h (e.g., *Zfand4, Tox2*, and *Slc14a2*) in the zebrafish and to 48 h (e.g., *Deg2, Ogfod1*, and *Ifitm1*) in the mouse. Gene transcription was apparently not prevented due to a lack of energy or resources. Several genes exhibited apparent regulation by feedback loops in their transcriptional profiles (e.g., *Rbm45* and *Cdc42* genes in the mouse (Fig 8 and 9, respectively; (Fig S3)). Hence, an underlying regulatory network appears to be still turning “on” and “off’ genes in organismal death.

### Interrupt the shutdown?

A living biological system is a product of natural selection and self-organizing processes [196]. Genes are transcribed and proteins translated in response to genetic and epigenetic regulatory networks that sustain life. In organismal death, we assumed most of the genetic and epigenetic regulatory networks operating in life would become disengaged from the rest of the organism. However, we found that “dead” organisms turn genes on and off in a non-random manner (Fig 2D and 2E). There is a range of times in which genes are upregulated and transcript abundances are maximized. While most genes are upregulated within 0.5 h postmortem (Fig 11), some are upregulated at 24 h and still others at 48 h. A similar pattern occurs with peak transcript abundances and the timing when upregulation is apparently stopped. These differences in timings and abundances suggest some sort of global regulation network is still operating in both organisms. What makes gene expression of life different from gene expression in death is that postmortem upregulation of genes offers no obvious benefit to an organism. We argue that self-organizing processes driven by thermodynamics are responsible for the postmortem upregulation of genes. We emphasize that such postmortem conditions could allow investigators to tease apart evolution from self-organizing processes that are typically entangled in life.

Since our results show that the system has not reached equilibrium yet, it would be interesting to address the following question: *what would happen if we arrested the process of dying by providing nutrients and oxygen to tissues?* It might be possible for cells to revert back to life or take some interesting path to differentiating into something new or lose differentiation altogether, such as in cancer. We speculate that the recovering cells will likely depend on the postmortem time – at least when such potentially interesting effects might be seen.

### Methodological validity

The Gene Meter approach is pertinent to the quality of the microarray output obtained in this study because conventional DNA microarrays yield noisy data [197,198]. The Gene Meter approach determines the behavior of every microarray probe by calibration – which is analogous to calibrating a pH meter with buffers. Without calibration, the precision and accuracy of a meter is not known, nor can one know how well the experimental data fits to the calibration (i.e., *R^2^*). In the Gene Meter approach, the response of a probe (i.e., its behavior in a dilution series) is fitted to either Freundlich or Langmuir adsorption model, probe-specific parameters are calculated. The “noisy” or “insensitive” probes are identified and removed from further analyses. Probes that sufficiently fit the model are retained and later used to calculate the abundance of a specific gene in a biological sample. The models take into consideration the non-linearity of the microarray signal and the calibrated probes do not require normalization procedures to compare biological samples. In contrast, conventional DNA microarray approaches are biased because different normalizations can yield up to 20 to 30% differences in the up- or down-regulation depending on the procedure selected [199–202]. We recognize that next-generation-sequencing (NGS) approaches could have been used to monitor gene expression in this study. However, the same problems of normalization and reproducibility (mentioned above) are pertinent to NGS technology [203]. Hence, the Gene Meter approach is currently the most advantageous to study postmortem gene expression in a high throughput manner.

### Practical implications

The postmortem upregulation of genes in the mouse has relevance to transplantation research. We observed clear qualitative and quantitative differences between two organs (liver and brain) in the mouse in their degradation profiles (Fig 1). We also showed the upregulation of immunity, inflammation and cancer genes within 1 h of death (Fig 11). It would be interesting to explore if these differences are comparable to what occurs in humans, and we wonder how much of the transplant success could be attributed to differences in the synchronicity of postmortem expression profiles rather than immunosuppression agents [204,205]. Our study provides an alternative perspective to the fate of transplant recipients due to the upregulation of regulatory and response genes, after the sample has been harvested from the donor.

## Conclusion

This is the first study to demonstrate active, long-term expression of genes in organismal death that raises interesting questions relative to transplantology, inflammation, cancer, evolution, and molecular biology.

## Authors’ contributions

DT, RN, TDL and AEP designed the study. AEP, RN, TDL carried out the molecular genetic studies. BGL and AEP determined the statistically significant upregulated genes. SS and PAN annotated the genes. AEP and PAN conducted the statistical analyses and wrote the manuscript. All authors read and approved the final manuscript.

## Acknowledgements

We thank Till Sckerl for help with sacrificing the mice, Nicole Thomsen for help with tissue and RNA processing, and Elke Blohm-Sievers for microarray work. We thank Russell Bush for his helpful advice on thermodynamics and entropic barriers and M. Colby Hunter for proof-reading the manuscript.

## Additional Files

**This PDF file includes:**

Supplementary Text

Figs S1, S2 and S3

Table S1

**Other Additional Files for this manuscript include the following:**

Data files S1 to S8 as zipped archives:

File S1. MiceProbesParameters.txt

File S2. FishProbesParameters.txt

File S3. Mouse_liver_Log10_AllProfiles.txt

File S4. Mouse_brain_Log10_AllProfiles.txt

File S5. Fish_Log10_AllProfiles.txt

File S6. MiceProbesSeq.txt

File S7. FishProbesSeq.txt

File S8. Gene annotation lit refs v3.xls

## SUPPLEMENTARY TEXT

Fig S1. Bioanalyzer results showing total mRNA from the zebrafish.

Fig S2. Bioanalyzer results showing total mRNA from the mouse.

Fig S3. Transcriptional profiles in the zebrafish (*Acer3* gene) and mouse (*Cdc42* and *Rbm45* genes) by postmortem time.

Table S1. Total mRNA extracted (ng/μl tissue extract) from zebrafish by time and replicate sample.

Table S2. Total mRNA extracted (ng/μl tissue extract) from mouse organ/tissue by time and replicate sample.

Table S3. % of global regulator genes and response genes.

**Fig S1.**
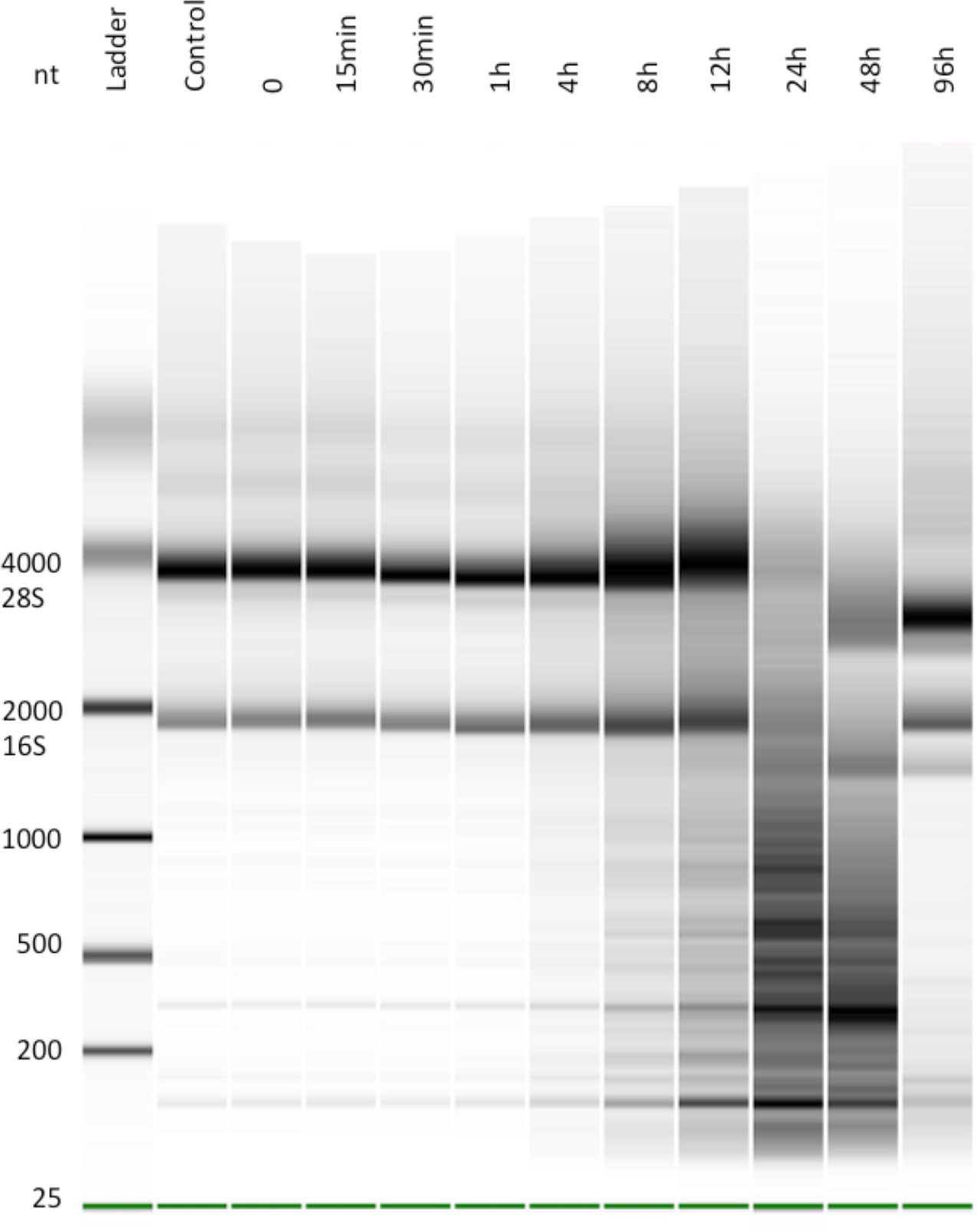
Bioanalyzer results showing total mRNA from the zebrafish. Only one replicate per sampling time is shown. The dominant bands represent the 28S and 18S rRNAs.

**Fig S2.**
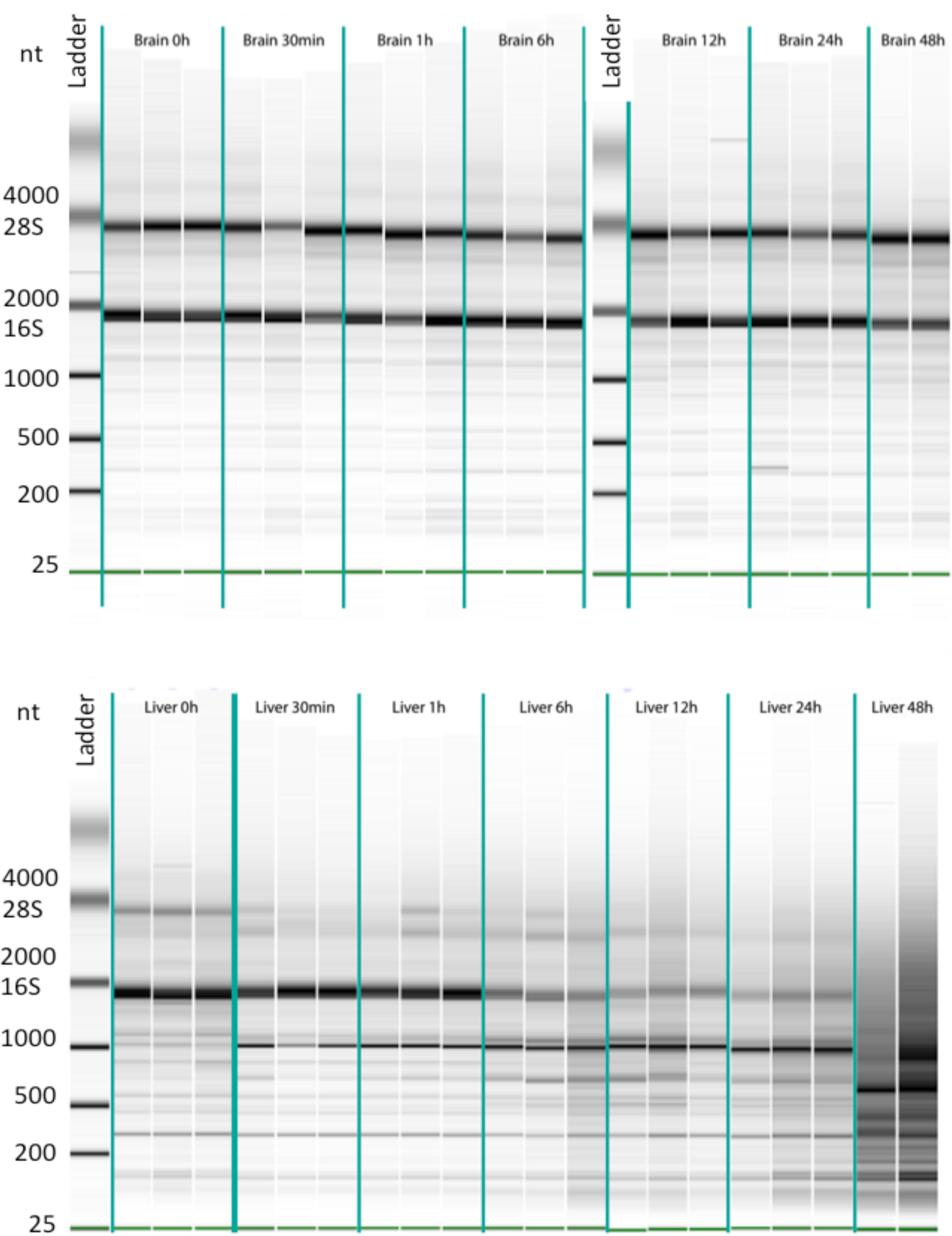
Bioanalyzer results showing total mRNA from the mouse. All replicates per sampling time are shown. The dominant bands represent the 28S and 18S rRNAs.

**Fig S3.**
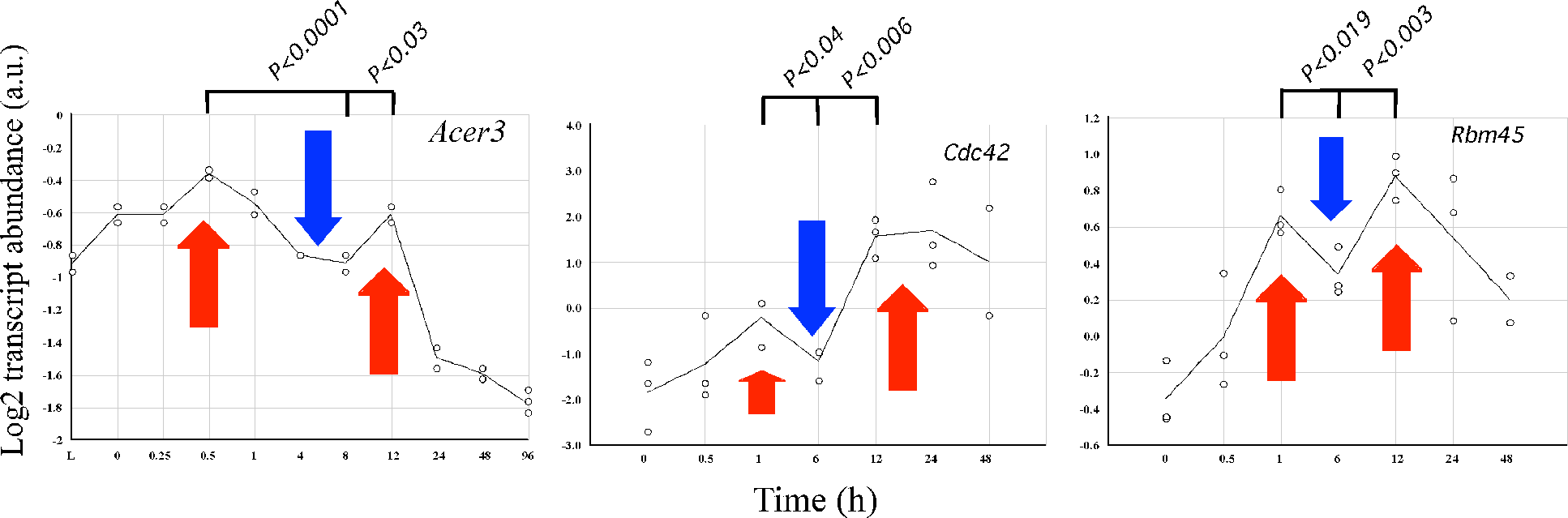
Transcriptional profiles in the zebrafish (*Acer3* gene) and mouse (*Cdc42* and *Rbm45* genes) by postmortem time. Red arrows, up-regulation; blue arrows, down-regulation. One-way T-tests show significant differences between means. Results suggest that the differences in up- and down regulation by postmortem time are due to changes in regulation rather than changes in “residual transcription levels”.

**Table S1.** Total mRNA extracted (ng/μ1 tissue extract) from zebrafish by time and replicate sample.

| Time (h) | Repl#1 | Repl#2 |
| --- | --- | --- |
| Live | 1606 | 1648 |
| 0 | 1632 | 1481 |
| 0.25 | 1768 | 1573 |
| 0.5 | 1457 | 1481 |
| 1 | 1864 | 1514 |
| 4 | 1251 | 1651 |
| 8 | 1605 | 1499 |
| 12 | 1365 | 1422 |
| 24 | 1087 | 539 |
| 48 | 339 | 428 |
| 96 | 183 | 183 |

a, no replicate taken

**Table S2.** Total mRNA extracted (ng/μ1 tissue extract) from mouse organ/tissue by time and replicate sample.

| Organ | Time (h) | Repl#1 | Repl#2 | Repl#3 |
| --- | --- | --- | --- | --- |
| Liver | 0 | 432 | 387 | 443 |
| Liver | 0.5 | 533 | 651 | 569 |
| Liver | 1 | 421 | 426 | 601 |
| Liver | 6 | 541 | 624 | 528 |
| Liver | 12 | 839 | 450 | 845 |
| Liver | 24 | 1021 | 510 | 1066 |
| Liver | 48 | 1453 | 1197 | -<sup>a</sup> |
| Brain | 0 | 169 | 226 | 210 |
| Brain | 0.5 | 174 | 166 | 410 |
| Brain | 1 | 194 | 485 | 264 |
| Brain | 6 | 401 | 269 | 256 |
| Brain | 12 | 379 | 258 | 203 |
| Brain | 24 | 249 | 324 | 400 |
| Brain | 48 | 397 | 310 | - |

a, no replicate taken

**Table S3.**
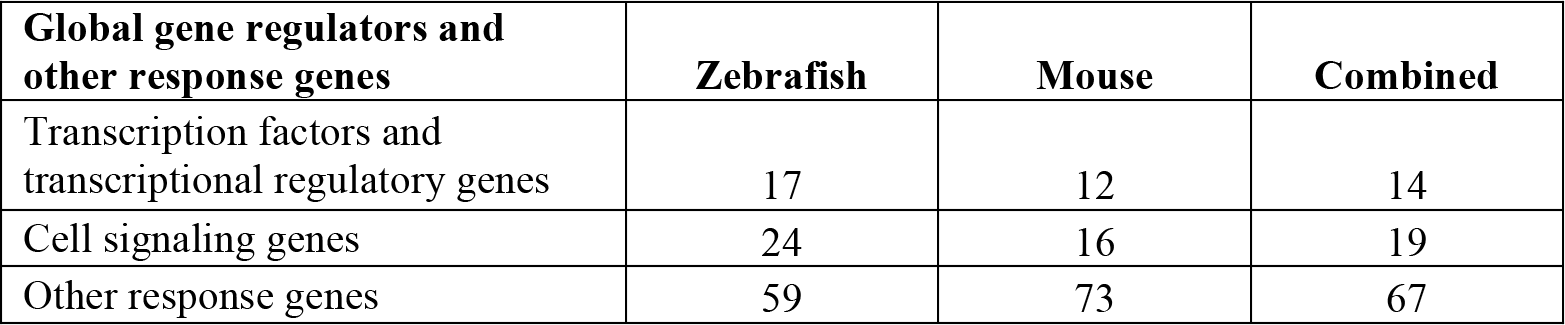
% of global regulator genes and response genes. Approx. 33% of all upregulated genes are involved in global regulation.

| **Global gene regulators and other response genes** | **Zebrafish** | **Mouse** | **Combined** |
| --- | --- | --- | --- |
| Transcription factors and transcriptional regulatory genes | 17 | 12 | 14 |
| Cell signaling genes | 24 | 16 | 19 |
| Other response genes | 59 | 73 | 67 |

